# Genomic and transcriptomic analyses of the elite rice variety Huizhan provide insight into disease resistance and heat tolerance

**DOI:** 10.1101/2024.03.30.587414

**Authors:** Wei Yang, Zhou Yang, Lei Yang, Zheng Li, Zhaowu Zhang, Tong Wei, Renliang Huang, Guotian Li

## Abstract

Rice is an important crop and serves as a model for crop genomics and breeding studies. Here, we used Oxford Nanopore ultra-long sequencing and next-generation sequencing technologies to generate a chromosome-scale genome assembly of Huizhan, a disease-resistant and heat-tolerant *indica* rice variety. The final genome assembly was 395.20 Mb with a scaffold N50 of 31.87 Mb. We identified expanded gene families in Huizhan that are potentially associated with both organ growth and development, as well as stress responses. We observed that three functional rice blast resistance genes, including *Pi2*, *Pia* and *Ptr*, and bacterial blight resistance gene *Xa27*, likely contribute to disease resistance of Huizhan. In addition, integrated genomics and transcriptomics analyses show that *OsHIRP1*, *OsbZIP60*, the SOD gene family, and various transcription factors are likely involved in heat tolerance of Huizhan. Results presented in this study will serve as a valuable resource for rice functional genomics studies and breeding.

## Introduction

Rice (*Oryza sativa*) is a major staple crop that feeds more than half of the world’s population (Zeng et al., 2017). Cultivated rice comprises two major subspecies, the *Geng*/*Japonica* (GJ) group and the *Xian*/*Indica* (XI) group (Wang et al., 2018). As a leading model for crop plants, the availability of rice genomes for rice varieties including Nipponbare, 93-11 (Qin et al., 2021), Shuhui498 (R498) (Du et al., 2017), Zhenshan 97 (ZS97), and Minghui 63 (MH63) (Song et al., 2021), has significantly facilitated functional genomics studies and provides valuable resources for rice breeding. The XI rice accessions have been reported to have higher genetic diversity than GJ rice accessions (Song et al., 2021). Therefore, establishing additional high-quality genomic resources for XI rice varieties is necessary to harness these genetic varieties in basic studies and rice breeding.

Global climate warming poses a significant threat to rice production, with extreme temperature stress being a particular concern (Kim et al., 2019). Heat stress significantly decreases the viability of rice pollens during the reproductive stage, resulting in a 10% reduction in grain yield per 1 °C increase in the dry season (Peng et al., 2004) based on field experiments. Moreover, elevated temperatures negatively impact rice grain quality, leading to an increased chalky appearance (Xu et al., 2020). Developing heat-tolerant varieties has often relied on overexpressing genes related to heat tolerance, such as *Thermo-tolerance 3* (*TT3*) in rice (Zhang et al., 2022). However, the widespread adoption of this strategy in rice production has been limited by the specificity of overexpressed genes related to heat tolerance or the intricate interaction of various abiotic stressors (El-Esawi and Alayafi, 2019). Hence, exploring rice varieties with exceptional heat tolerance and desirable agricultural traits could offer valuable insight into enhancing rice heat tolerance.

Plant diseases pose another severe threat to global food security, leading to substantial losses in agricultural productivity and quality (Chen et al., 2019). For example, rice blast caused by the fungal pathogen, *Magnaporthe oryzae*, results in yield losses equivalent to feeding over 60 million people worldwide each year (Sakulkoo et al., 2018). Plants initiate pattern-triggered immunity (PTI) by recognizing microbial or pathogen-associated molecular patterns (MAMPs or PAMPs), such as flagellin (Zipfel and Felix, 2005), through pattern recognition receptors (PRRs) (Jones and Dangl, 2006). On the other hand, effector-triggered immunity (ETI) is an intracellular response initiated when nucleotide binding site-leucine rich repeat (NBS-LRR) proteins or *R* genes recognize specific pathogen effectors. In rice, the *Pi* gene confers resistance to rice blast. *Pi* genes typically encode NBS-LRR proteins, except for *Pi21*, *Pid2* and *Ptr*. Until now, dozens of *R* genes have been cloned, such as *Pita*, *Pi37*, *Pit*, *Pish*, *Pi64*, *Pi2*, and *Ptr* (Xiao et al., 2020). However, due to the high variability and rapid evolution of the pathogen population, the resistance conferred by a single *R* gene often becomes ineffective within a few years. By pyramiding appropriate *R* genes/alleles, it is possible to effectively achieve durable resistance against rice blast, such as *Pik* + *Pita*, *Pi9* + *Pi54* (Wu et al., 2015), and *Pi2* + *Pib* (Wang et al., 2021). As a result, breeders tend to favor selecting rice varieties that possess multiple *R* genes, enabling durable and broad-spectrum resistance (Li et al., 2019).

Rice is susceptible to a wide range of biotic and abiotic stresses. Previous research has shown that the *Xa7* gene, renowned for its resistance to bacterial leaf blight, exhibits superior efficacy during periods of heat stress. Conversely, other *R* genes associated with bacterial leaf blight, such as *Xa21*, show decreased efficacy at elevated temperatures (Cohen and Leach, 2020). Furthermore, rice exhibits reduced resistance to rice blast under heat stress, thereby expediting pathogen colonization of rice tissues and exacerbating disease severity (Onaga et al., 2017). Consequently, the availability of various genomic resources targeting combined stress conditions can significantly enhance our comprehension of the rice-pathogen-high temperature model and facilitate precise breeding efforts (Cohen and Leach, 2020).

Huizhan, a recently developed XI rice variety is known for its robust disease resistance, heat tolerance and high yield, with consistent performance across various environments, and stable essential characteristics. Furthermore, Huizhan serves as a premium parental line for the improvement of hybrid rice, as demonstrated by the hybrid rice cultivar Huiliangyouhuizhan, with Huizhan as the pollen donor. This variety has gained widespread adoption in the mid-lower reaches of the Yangtze River in China.

To facilitate the genetic analysis of Huizhan and apply it to precision breeding, we have *de novo* assembled a high-quality, chromosome-level genome of Huizhan using Oxford Nanopore ultra-long reads and next-generation sequencing short reads. We identified 39,558 protein-coding genes in Huizhan and detected small genomic variations by comparing its genome with those of other rice varieties. We found that three rice blast resistance genes, including *Pi2*, *Pia* and *Ptr*, as well as the *Xa27* gene, for bacterial blight resistance, remain complete and unmutated in Huizhan. Compared to 16 other rice varieties, gene families related to organ growth and development, lipid transport and positive signaling transduction have significantly expanded and evolved. By comparing the leaf and flower transcriptomes of two rice varieties Huizhan and Zhonghua 11 (ZH11), we identified the differentially expressed genes (DEGs) and provide insights into the common and distinct responses of Huizhan and ZH11 to heat stress. We have identified eight up-regulated genes associated with heat tolerance. Based on previous research on thermo-perception and thermo-responses in rice, we identified key genes with the focus on heat-response and transcription factors, including *OsbZIP60*, the SOD gene family, and HSFs, which likely play a critical role in heat tolerance of Huizhan, gaining insights into the possible genetic basis underlying heat tolerance. The high-quality Huizhan genome facilitates rice functional genomics and precision breeding for disease resistance and heat tolerance.

## Materials and methods

### Plant materials

The *Oryza sativa* accession Huizhan was obtained from Jiangxi Academy of Agricultural Sciences. The seeds were planted in a growth chamber at the experimental station of Huazhong Agricultural University (Wuhan, China) in May. The light: dark period was around 13/11 h at 28 °C. Fresh young leaf tissues from 4-week-old seedlings were used in sequencing. Rice for RNA sequencing were planted in the Nanchang region, Jiangxi Province, China (28°33’42”E, 115°56’5’’N), sown on 10 May 2022 and transplanted on 1 August to ensure high temperature conditions. Fresh samples were washed and preserved in liquid nitrogen for RNA sequencing.

### Library construction and sequencing

The DNA sample was isolated using the cetyltrimethylammonium bromide (CTAB) method. To assess the quality and quantity of DNA preparations, a NanoDrop One UV–Vis spectrophotometer (Thermo Fisher Scientific, USA) and Qubit 3.0 Fluorometer (Invitrogen, USA) were utilized, respectively. Selected high-quality DNA was used for Oxford Nanopore sequencing and short reads sequencing. For the ultra-long reads library, genomic DNA (>50 kb) was selected with the SageHLS HMW library system and then processed with the Ligation Sequencing 1D kit (SQK-LSK109) (Wang et al., 2022). The resulting libraries were sequenced on the Promethion platform (Oxford Nanopore Technologies, UK) at Grandomics’ Genome Center (Wuhan, China). Using Guppy v5.0.11 (Wick et al., 2019) for rapid base calling, we obtained a total of 808,518 ultra-long reads with an N50 of 53,290 bp. For short reads sequencing, a 150-bp paired-end reads library was constructed and sequenced using the MGIEasy RNA Library Prep Set on the DNBSEQ-T7 platform. Low quality reads were filtered by Fastp v0.23.2 (Chen et al., 2018) with default parameters.

### RNA preparation and sequencing

RNA from Huizhan was extracted with Trizol. To improve the quality of genome annotation, fresh tissues including roots from 2-week-old seedlings, leaves from 4-week-old seedlings and flag leaf, shoots from 6-week-old seedlings, panicles, ears (2-3 days before and 2-3 days after heading), flowers were collected for RNA sequencing on the DNBSEQ-T7 platform. For transcriptome assays of heat-stressed plants, leaf and flower samples from three biological replicates of Huizhan and ZH11 were collected and sequenced on the NovaSeq 6000 platform (Illumina) following the manufacturers’ recommended procedures.

### *De novo* genome assembly and evaluation

Jellyfish v2.2.10 (Marcais and Kingsford, 2011) and Genomescope v2.0 (Ranallo-Benavidez et al., 2020) were used for *k*-mer analysis. The 19-mer frequency distributions were used to estimate the genome size of the Huizhan in all clean short reads. The ultra-long reads have a tremendous advantage in the challenges of genome gaps and highly repetitive regions (Deng et al., 2022; Song et al., 2021). Raw ultra-long reads of the Huizhan genome were filtered by removing reads with a q-value of less than 7. NextDenovo v2.5.0 (https://github.com/Nextomics/NextDenovo) was used to *De novo* assembly of clean ultra-long reads. To enhance assembly accuracy, the resulting assembly from NextDenovo was polished using Nextpolish v1.4.0 (parameters: task = best) with the ultra-long reads and short reads (Hu et al., 2020). Then, the contigs were anchored to the scaffold/chromosome level using RaGOO v1.1 with reference genome MH63 (Alonge et al., 2019).

The assembly continuity of the Huizhan genome was assessed by the long terminal repeat (LTR) assembly index (LAI) value, which identified LTR retrotransposons from the outputs of the LTR_retriever v2.9.0 (Ou and Jiang, 2018) pipeline. We evaluated the completeness of the assembled genome with the BUSCO v5.3.2 (Simao et al., 2015) with embryophyta standard database 10. We also assessed assembly consensus quality (QV) and completeness measurements based on *k*-mer using Merqury v1.3 (Rhie et al., 2020). The average depth of coverage analysis with mapped short reads and ultra-long reads to the Huizhan genome was performed using bwa v0.7.17-r1188 (https://github.com/lh3/bwa) and minimap2 v2.26-r1175 (Li, 2018).

### *CentO*-enriched regions and telomere sequences identification

We built databases from the *CentO* satellite repeat sequences in rice and searched the Huizhan genome with BLASTN v2.14.0 (parameters: -evalue 1e-10) (Altschul et al., 1990). BEDtools v2.31.0 (Quinlan, 2014) was subsequently used to merge the resulting alignments (parameters: -d 50000). The region that exceeded 10 kb in length and possessed the highest number of *CentO* repeats was identified as approximate *CentO*-enriched regions as described. Telomere sequences were searched in the Huizhan genome with quarTeT v1.1.6 (Lin et al., 2023).

### Repeat annotation

Dfam v3.3 (Storer et al., 2021) and RepBase-20181026 databases (http://www.girinst.org/) were used to construct a homology-based repeat library. To further investigate repetitive elements, a *de novo* LTR library was generated using the EDTA v2.0.1 pipeline (Ou et al., 2019). Unclassified LTR elements were analyzed with the DeepTE (Yan et al., 2020). RepeatMasker v4.1.2 (http://www.repeatmasker.org) was used to identify the repetitive sequences in the Huizhan genome.

### Gene prediction and functional annotation

Gene prediction was based on integrated homology-based searches, evidence-based from RNA-seq data, and ab initio predictions. We used homology-based searches with GeMoMa v1.9 (Keilwagen et al., 2018) of the proteomes of rice (ZS97, MH63, 93-11, R498) (Qin et al., 2021; Song et al., 2021), maize (Mo-17) (Sun et al., 2018) and *Arabidopsis* (TAIR10) as hints. The homology-based gene models were subsequently combined and filtered gene models of low quality or redundancy were eliminated. Evidence-based gene models were generated by mapping to the RNA-seq dataset with Hisat2 v2.2.1 (Kim et al., 2015a). The potential transcripts were identified by a genome-guided approach using StringTie v2.2.1 (Pertea et al., 2015). We also used the *de novo* assembler Trinity v2.13.2 (Haas et al., 2013) to generate RNA-seq assemblies and then mapped them back to the Huizhan genome with PASApipeline v2.5.3 (Haas et al., 2008) (PASA-Trinity-set). Transdecoder v5.7.1 (available at https://github.com/TransDecoder/TransDecoder) was used to locate potential open reading frames. For ab initio annotation, GeneMark-ETP and Augustus v3.5.0 were trained using Braker v3.0.3 (Hoff et al., 2019) on the repeat-masked genome sequences. The transcriptomics data from multiple tissues and the

Eukaryota protein database from OrthoDB v11 (Kuznetsov et al., 2023) were utilized as inputs for Braker. SNAP (2006-07-28) (Korf, 2004) also predicted the de novo gene models based on PASA-Trinity-set. Gene models generated from all methods were joined by EVidenceModeler v2.1.0 (Haas et al., 2008). In addition, we performed two rounds of updates using PASApipeline v2.5.3 (Haas et al., 2008) with the default parameters. The annotation completeness of the Huizhan genome was assessed using BUSCO v5.3.2 (Simao et al., 2015) with embryophyta standard database 10.

Transfer RNAs (tRNAs) were identified with tRNAscan-SE v2.0.12 (Chan et al., 2021), and ribosomal RNAs (rRNAs) were predicted with RNAmmer v1.2 (Lagesen et al., 2007). Non-coding RNAs were identified using Rfam v14.9 and Infernal v1.1.4 (Nawrocki and Eddy, 2013).

The functional annotation of protein-coding genes was performed using three tools. Interproscan v5.63-95.0 (Jones et al., 2014) and eggNOG-mapper v2.1.12 (Huerta-Cepas et al., 2019) were used to obtain functional domains by searching against publicly available databases. Further, SwissProt, NR and KEGG databases were used for annotation using BMKCloud (https://www.biocloud.net/).

### Chloroplast genome assembly and annotation of Huizhan

The chloroplast genome of Huizhan was assembled using GetOrganelle v1.7.6.1 pipeline with the optimized parameters ‘-k 21,55,85,115,127 -F embplant_pt’ (Jin et al., 2020). The final chloroplast genome was annotated with the online software CPGAVAS2 (Shi et al., 2019) and GeSeq (Tillich et al., 2017). The complete chloroplast genomes of *O. sativa* XI group IR8 (GenBank: KM103367) and TN1 (GenBank: KM103368) served as the reference database (Kim et al., 2015b). Geneious v2023.1 (Kearse et al., 2012) was used to compare the annotation results and manually correct and complement problematic annotations. Finally, the assembly and annotation results were used to generate a circular map with Chloroplot (Zheng et al., 2020). Tandem Repeats Finder v4.09 (Benson, 1999) was used to find tandem repeats and REPuter (Kurtz et al., 2001) was used to determine forward, palindromic, reverse, and complement repeats. Simple sequence repeats (SSRs) were identified with Misa-web (Beier et al., 2017). Codon preference analyses were performed using EMBOSS (Rice et al., 2000).

### Collinearity analysis of different accessions

The six genomes of the XI rice varieties and the GJ rice variety LTH genome were aligned to reference genome Huizhan with Nucmmer v4.0.0beta2 (Marcais et al., 2018) with parameters settings ‘-g 1000 -c 90 -l 40’. The resulting alignment blocks were filtered using Delta-filter with the parameters ‘-r -q’ to obtain one-to-one alignment results.

### SNPs and InDels analysis

SNPs and InDels were identified from the resulting filtered delta files with Show-snps with parameters settings ‘-ClrTH’. ANNOVAR (Wang et al., 2010) was employed for the annotation of the identified SNPs and InDels.

### Identification of PAVs

Show-diff in Mummer v4.0.0beta2 (Marcais et al., 2018) was used to select for the potential PAV sequences (Song et al., 2020). Sequences with feature type BRK and DUP were excluded, and only sequences with a minimum length of 100 bp were retained. Further, minimap2 (Li, 2018) was used to map the candidate PAV sequences to the LTH genome with the parameter setting ‘-x asm10’, and the sequences covering >80% were filtered out to obtain the final PAV regions. Genes that exhibited an overlap of more than 80% with the PAV regions were identified as PAV-related genes (Song et al., 2020). To identify enrichment results of PAV-related genes, we used the R package ‘clusterProfiler’ v4.10.0 (Yu et al., 2012) to construct a package based on the annotation results from eggNOG-mapper tools (Huerta-Cepas et al., 2019).

### Analysis of *Pi* and *Xa* genes

We selected a total of 39 published the *Pi* proteins for our analysis, which included *Pi1-5*, *Pi1-6*, *Pi2*, *Pi21*, *Pi36*, *Pi37*, *Pi5-1*, *Pi5-2*, *Pi50_NBS4_3*, *Pi50_NBS4_1*, *Pi7-1*, *Pi7-2*, *Pi9*, *Pia*, *Pib*, *Pigm-R6*, *Pii-1*, *Pii-2*, *Pik-h*, *Pikp-1*, *Pikp-2*, *Pik-1*, *Pik-2*, *Pikm1*, *Pikm2*, *Piks-1*, *Piks-2*, *Pish*, *Pit*, *Pita*, *Piz-t*, *Pizh*, *Ptr*, *OsRGA5*, *Pid3*, *Pi63*, *Pi56*, *Pi54, Pb1* (Yang et al., 2022).The 16 *Xa* genes also included *Xa1*, *Xa2/Xa31*, *Xa3*, *Xa26*, *Xa4*, *Xa5*, *Xa7*, *Xa10*, *Xa13*, *Xa14*, *Xa21*, *Xa23*, *Xa25*, *Xa27*, *Xa41*, and *Xa45*. We used the BLASTP (Altschul et al., 1990) to align the selected proteins with the protein set of the Huizhan genome. Hits with an identity < 75% and an E-value > 1e-10 were filtered out.

### Phylogenetic analysis

OrthoFinder v2.5.5 (Emms and Kelly, 2015) was used to construct gene families in *O. brachyantha*, *O. punctata*, *O. glaberrima*, *O. rufipogon*, Nipponbare, ZH11, Kitaake, Huizhan, ZS97, MH63, FH838, R498, G46, 93-11, IR64, TM, and Tumba (Qin et al., 2021). Using MAFFT v7.475 (Katoh and Standley, 2013), we independently aligned the protein sequences of 7,137 single-copy genes from 17 species and concatenated them into a superalignment matrix. Maximum likelihood trees of the 17 species were constructed by the maximum likelihood (ML) method using RAxML v8.2.12 (Stamatakis, 2014) with 100 bootstrap replicates. The calibration of species divergence time was conducted using MCMCTree in Phylogenetic Analysis by Maximum Likelihood (PAML) v4.9j (Yang, 2007) based on four-fold degenerate sites (4DTV) sequences in single-copy orthologous genes and the TimeTree website (Kumar et al., 2022). The CAFÉ5 program (De Bie et al., 2006) was employed to analyze the gene families expansion and contraction with the default parameters. ITOL website (Letunic and Bork, 2007) was used to visualize of the phylogenetic tree. GO enrichment analyses of the genes in the expanded families and the positive selective genes were performed with the R package ‘clusterProfiler’ v4.10.0 (Yu et al., 2012) based on the annotation results from InterProScan (Jones et al., 2014).

### Quantification and identification of DEGs

We used the Fastp v0.23.2 (Chen et al., 2018) to filter the low-quality RNA sequencing reads and aligned the filtered clean reads to the reference genome Nipponbare from Rice Genome Annotation Project Release 7 (RGAP) (Kawahara et al., 2013) using Hisat2 v2.2.1 (Kim et al., 2015a). Transcripts Per Million (TPM) of each gene was quantified and calculated using FeatureCounts from the R package Rsubread v2.12.0 (Liao et al., 2019). Differentially expressed genes (DEGs) were analyzed with DESeq2 v1.38.0 (Love et al., 2014) with |log2(fold change)| > 1 and *P*-adj < 0.05. Gene ontology (GO) and KEGG pathway (Kanehisa and Goto, 2000) enrichment analysis of the DEGs were performed using Rice Gene Index (https://riceome.hzau.edu.cn/) (Yu et al., 2023) and KOBAS v3.0 (Bu et al., 2021). Heat stress responsive related genes were obtained from the funRiceGenes database (Yao et al., 2018). The expression correlation of the different transcription factors was determined with Pearson correlation analysis (*p*-value < 0.05).

### Pollen viability examination

To compare male sterility at high temperatures, we focused on pollen grain viability of anthers of Huizhan and ZH11. During the flowering period on 25th August 2022 at Nanchang region, Jiangxi Province, China (28°33’42’’E, 115°56’5’N), we collected three representative spikes of each cultivar at the full bloom period. Three anthers were removed with tweezers from each flower and then immersed in 20 μl of Lugol′s solution. The anthers were dehisced with tweezers on a slide, and images were taken under a microscope (DMI3000 B, Leica, Germany) after removing anther wall debris. The number of observed pollen grains was counted from the digital images using ImageJ software (http://imagej.nih.gov/ij/). The pollen viability was calculated from the ratio of the number of intensely colored pollen grains to the total number of pollen grains. Each cultivar was evaluated 10 times.

## Results

### Sequencing, assembly and assessment of the Huizhan genome

To assemble the genome of Huizhan in high-quality (Figure 1a), we generated ∼31.6 Gb ultra-long reads (91× coverage) with an N50 size of 53,290 bp and ∼59.4 Gb filtered short reads (149× coverage) (Table S1). Furthermore, we generated ∼69.7 Gb RNA-seq data from multiple tissue types of Huizhan at different developmental stages (Table S2). Prior to the *de novo* assembly, the genome size was estimated at ∼396.11 Mb based on *k*-mer frequency analyze (Table 1, Figure S1). The assembly of the genome was accomplished through a hybrid strategy. The contigs based on ultra-long reads were *de novo* assembled using NextDenovo v2.5.0. We iteratively polished this draft assembly with ultra-long reads and short reads. The final assembly consists 15 contigs with a contig N50 of 31.35LMb (Table 1). Using RaGOO v1.1 (Alonge et al., 2019), these contigs were sorted and oriented into 12 chromosome-level pseudomolecules, and 9 chromosomes were covered by a single contig, while pseudomolecules 4, 10 and 11 contained only one gap each (Figure 1b). The final genome assembly of Huizhan is 395.20 Mb (scaffold N50, 31.87 Mb) (Figure 1b, Table S3), which covers 99.8% of the estimated genome (Table 1).

**Figure. 1.**
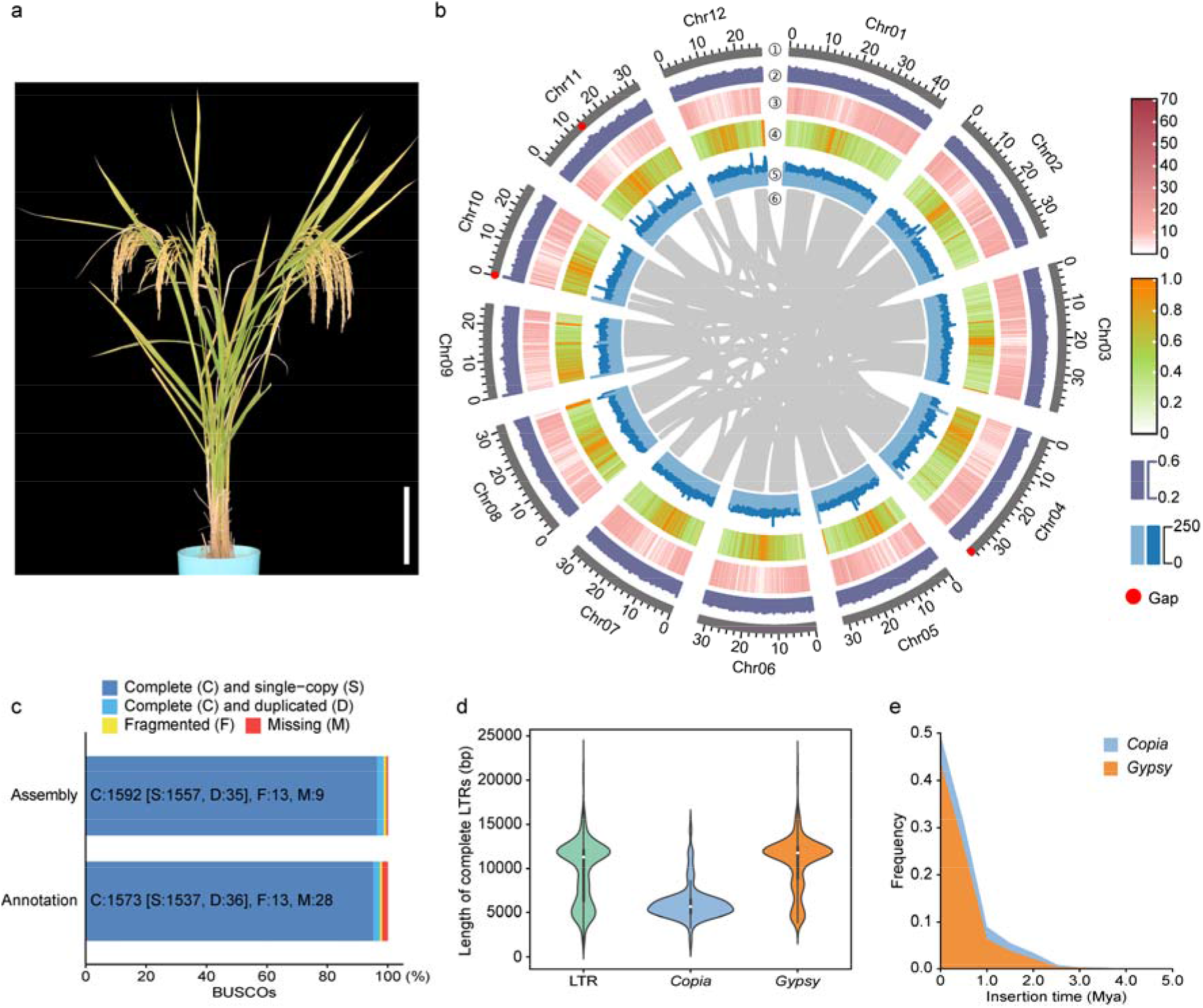
Genome of the *Oryza sativa* cultivar Huizhan. **a.** The rice cultivar Huizhan in the mature stage. Bar, 20Lcm. b. Circos plot of the Huizhan genome of 100-kb intervals. ①. The 12 chromosomes. The red dot indicates Gap. ②. GC content. ③. Gene density. ④. Repeat density. ⑤. The ultra-long reads and short reads coverage depth. ⑥. Syntenic blocks (3,461 gene pairs, 179 blocks). c. Evaluation of Huizhan genome assemblies and annotation using Benchmarking Universal Single-copy Orthologs (BUSCO) analysis. d. Length distribution of complete long terminal repeat retrotransposons (LTR-RTs) (whiskers and black boxes represent average length distribution, and white circles represent median). e. The insertion time (mya) distribution of intact LTR-RTs.

**Table. 1.** Genome assembly statistics of Huizhan.

|  | Huizhan | Zhenshan 97 | Minghui 63 |
| --- | --- | --- | --- |
| Assembly features |  |  |  |
| Estimated genome size (Mb) | 396.11 | - | - |
| Total size of assembled contigs (bp) | 395,196,687 | 391,561,630 | 395,765,488 |
| Number of contigs | 15 | - | - |
| Contig N50 (Mb) | 31.35 | - | - |
| Number of scaffolds/ pseudomolecules | 12 | 12 | 12 |
| Scaffold N50 (Mb) | 31.87 | 32.12 | 31.92 |
| GC content (%) | 43.65 | 43.62 | 43.65 |
| Ultra-long read-mapping rate (%) | 99.78 | - | - |
| Short read-mapping rate (%) | 99.91 | - | - |
| BUSCOs (%) | 98.7 | 99.88 | 99.88 |
| LTR assembly index | 24.22 | 24.01 | 22.74 |
| Repeat content (%) | 51.84 | 53.78 | 53.95 |
| Number of telomeres | 15 | 19 | 22 |
| Gene models |  |  |  |
| Number of predicted genes | 39,588 | 39,258 | 39,406 |
| Mean coding sequence length (bp) | 3,219.51 | 3256.97 | 3234.9 |
| Numbers of transcripts | 59,227 | 60,935 | 59,903 |
| Mean number of exons per gene | 4.37 | 4.8 | 4.13 |
| Mean exon length (bp) | 350.05 | 344.26 | 342.88 |

We identified *CentO*-enriched regions using the 155-165 bp *CentO* satellite repeat (Cheng et al., 2002) (Table S4). For the telomeric regions, we employed the criteria of having more than 100 seven-base telomeric repeats and identified a total of 15 telomeres, with Chr01, Chr03, Chr09, and Chr12 being telomere-to-telomere pseudomolecules (Figure S2, Table S5).

To estimate the accuracy of the assembly, both ultra-long reads and short reads were aligned to the genome, resulting in mapping ratios of 99.78% and 99.91%, respectively (Table 1, Figure S3). The completeness of genome was assessed by identifying 1,592 complete genes (98.7%) from embryophyta_odb10 databases using Benchmarking Universal Single-Copy Orthologs (BUSCO) v5.3.2 (Table 1, Figure 1c, Table S6). Furthermore, we used LTR retrotransposons (LTR-RTs) assessment to evaluate genomic continuity, which revealed an LTR Assembly Index (LAI) of 24.22, within the gold standard (≥ 20), for the Huizhan genome (Table 1) (Ou et al., 2018). We also used Merquery based on an efficient *k*-mer set with the clean short reads (Rhie et al., 2020), which indicated QV scores around 39.51 for 12 chromosomes (Table S7). In conclusion, these results collectively demonstrate the high-quality of the Huizhan genome.

### The chloroplast genome of Huizhan

The chloroplast is an important organelle that contributes to genetic diversity and stress responses (Kim et al., 2015b; Schwenkert et al., 2022; Zhang et al., 2022). The chloroplast genome of Huizhan was 134,501 bp in length assembled with GetOrganelle v1.7.6.1, with a GC content 39.01% (Figure S4). The chloroplast genome of Huizhan had a typical plastid genome structure and comprised two inverted repeat regions (IRA and IRB), a large single-copy (LSC) region and a small single-copy (SSC) region. A total of 134 chloroplast genes were annotated, including 85 protein-coding genes, 41 tRNAs and eight rRNAs. For the repeat elements in the chloroplast genome of Huizhan, we identified 26 tandem repeats, 23 forward repeats, 23 palindromic repeats, and four reverse repeats but not complement repeats. A total of 11 simple sequence repeats were identified in the chloroplast genome of Huizhan. Codon preference analyses showed that codon AAA, TTT and AAT are the most frequently used (Table S8).

### Repeats and genes annotation

The repetitive sequences of the Huizhan genome were identified by using a combined strategy of *de novo* and homology-based search. A total of 204.88 Mb (51.84% of assembled genome) repeat sequences were identified (Table 1, Table S9). The long terminal repeat (LTR) element family contains abundant types (27.37%), including *Gypsy* (23.32%), followed by *Copia* (3.64%). A *de novo* LTR library identified and classified a total of 2,786 intact LTR retrotransposons. The average lengths of complete LTR-RTs in *Copia* and *Gypsy* superfamilies were ∼L6.1 kb, and ∼L10.7 kb, respectively (Figure 1d), similar to the observations in the Nipponbare genome (Zhou et al., 2021). The frequency distribution of their insertion time exhibited a burst of LTR-RTs that occurred approximately 0.2 million years ago (mya), indicating that most of the retrotransposons were relatively recently inserted (Figure 1e). This observation provides evidence to support the notion that LTR-RTs have played an active role in driving the evolution of genomic diversification after intentional cultivation and domestication by humans (Carpentier et al., 2019).

To predict the protein-coding genes in Huizhan genome, we used EVidenceModeler (Haas et al., 2008), a pipeline that integrates homology-based searches, evidence from RNA-seq data and ab initio predictions, and predicted a total of 39,558 protein-coding genes with an average gene length of 3,219.51 bp (Table 1). The protein-level BUSCO score of the Huizhan genome was determined to be 97.4% (Figure 1c, Table S6). We functionally annotated these protein-coding genes using Interproscan (Jones et al., 2014), eggNOG-mapper (Huerta-Cepas et al., 2019) and BMKcloud, and found 38,157 (96.46%) genes (Table S10). Additionally, noncoding RNA sequences were annotated with homology-based methods. A total of 796 tRNAs, 1,485 rRNAs, 9,641 miRNAs, and 685 snRNAs were identified in the Huizhan genome.

### Genes families of Huizhan and 16 other *Oryza* species

To reveal the genetic relationship of Huizhan, we performed a comparative analysis between Huizhan and 16 other *Oryza* species, including *O. brachyantha*, *O. punctata*, *O. glaberrima*, *O. rufipogon*, Nipponbare, ZH11, Kitaake, ZS97, MH63, FH838, R498, G46, 9311, IR64, TM, and Tumba. We assigned 644,491 genes (96.2% of the total) to 77,147 orthogroups, including 52,803 orthogroups from all species, 1,278 species-specific orthogroups and 7,317 single-copy orthogroups (Figure 2a and 2c). To investigate the phylogenetic relationships between Huizhan and other *Oryza* species, we constructed a phylogenetic tree using the single-copy genes. The phylogenetic inference showed that Huizhan is the sister to a lineage composed of ZS97 and MH63, revealing a closer relationship between Huizhan and these two species (Figure 2d). Comparing Huizhan to ZS97, MH63 and Nipponbare, we identified 18,031 orthogroups shared by all four species in the 52,803 orthogroups, while 1,792 orthogroups were unique to the Huizhan genome (Figure 2b).

**Figure. 2.**
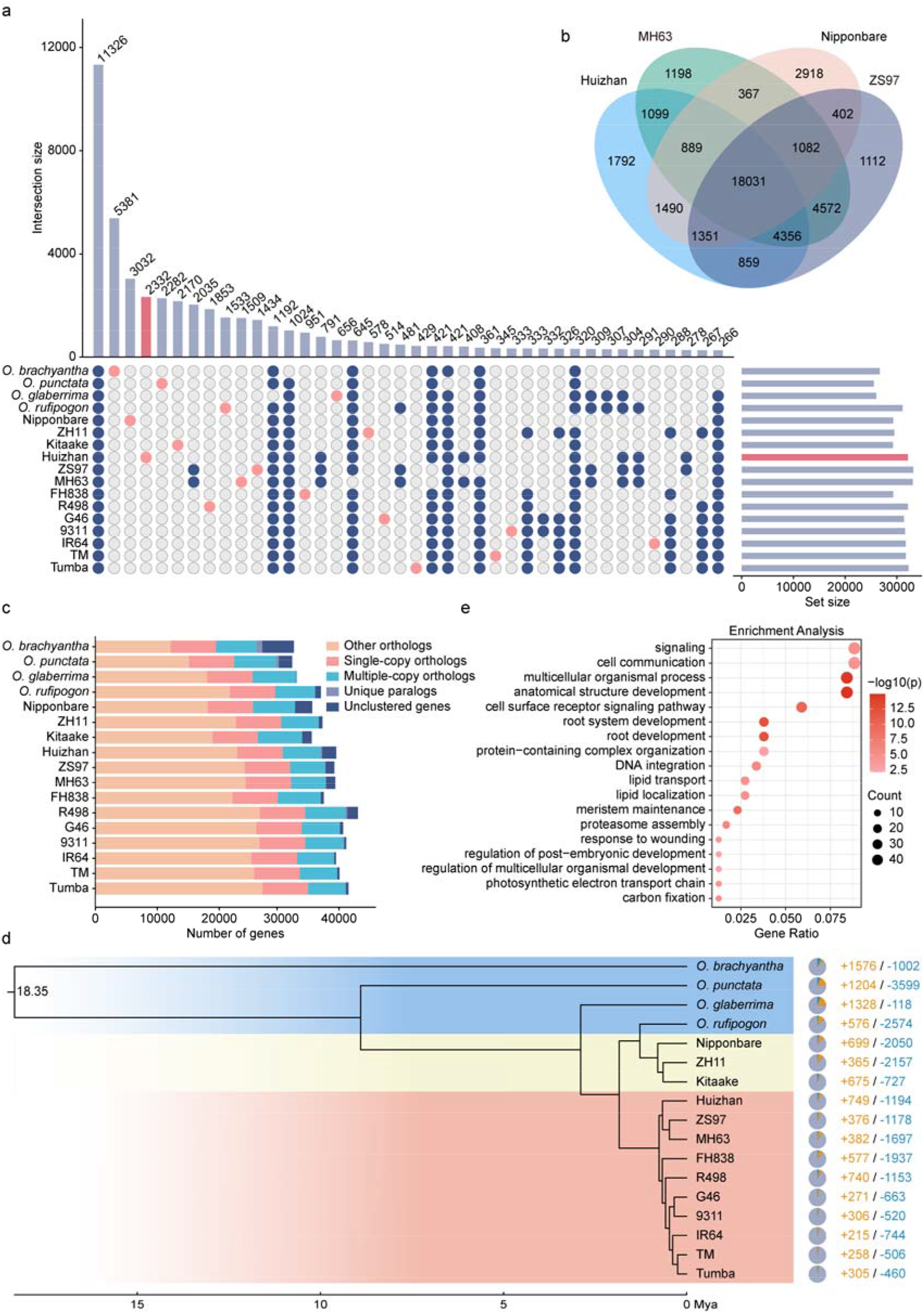
Genes family analysis between Huizhan and neighboring species. a. Upset plots show intersections of 77,147 orthogroups among 17 rice varieties. b. Venn diagram showing orthologous gene families among Huizhan, Nipponbare, ZS97, and MH63 genomes, excluding the orthogroups containing unassigned genes. c. Stacked bar charts showing numbers of orthologous genes found among 17 rice varieties. d. Phylogenomic analysis of Huizhan and 16 other rice varieties. The scale at the bottom indicates divergence times. The pie chart and the numbers on the right indicate gene family expansions (orange) and contractions (blue). e. GO enrichment results (biological process) of significantly expanded gene families of Huizhan.

The protein-coding genes in the Huizhan genome were classified into 2,332 unique families between Huizhan and 16 other *Oryza* species (Figure 2a), including 66 species-specific gene families and 2,266 non-clustered gene families. Gene ontology (GO) enrichment analysis showed that these non-clustered gene families were mainly associated with various biological processes, such as ‘positive regulation of developmental process’, ‘positive regulation of multicellular organismal process’, ‘organ growth’, and ‘regulation of organ growth’. We identified OsHZ15485 (*OsARGOS*) as the common gene among the four terms, which is closely related to cell proliferation and cell expansion (Wang et al., 2009).

Based on gene expansion and contraction analyses of Huizhan, we classified 749 and 1,194 gene families as expanded and contracted, respectively. The expanded genes were significantly enriched in Gene Ontology (GO) terms related to ‘cell communication’, ‘cell surface receptor signaling pathway’, ‘root development’, ‘lipid transport’, ‘response to wounding’, and ‘regulation of multicellular organismal development’ (Figure 2e).

### The global genome comparison between Huizhan and other rice varieties

We next performed a comparative analysis between the Huizhan genome and six other XI rice genomes. As expected, 91-95% of the Huizhan genome sequence was in one-to-one collinearity blocks with the six other genomes, indicating a high level of conservation among different XI rice varieties (Figure S5, Table S11). To further investigate genetic sequence variations, we identified ∼1.43 × 10^6^ single nucleotide polymorphisms (SNPs) and ∼0.30 × 10^6^ insertions and deletions (InDels) using the ‘Show-snps’ tool (Figure S6, Table S12). With the Huizhan genome as the reference, an average of 3.6 SNPs and 0.8 InDels were identified in per kb DNA (Figure S7). The SNPs and InDels were found to be more abundant in the intergenic regions and exhibited a positive correlation pattern throughout the Huizhan genome (Table S12). A chromosomal inversion is observed in the Huizhan genome at Chr12:0.75 Mb-1.10 Mb, distinguishing it from five other rice varieties, except ZS97 (Figure S8). This inverted region contains 53 genes, including WRKY transcription factors and non-specific lipid-transfer proteins (Table S13).

To evaluate disease resistance of Huizhan, we compared it to the widely used rice variety Lijiangxintuanheigu (LTH) (Figure 3a), which exhibits universal susceptibility to thousands of *Magnaporthe oryzae* isolates (Yang et al., 2022). The high-quality genome provided us a valuable resource for identifying possible presence-absence variations (PAVs). By comparing the two genomes, we identified 12,177 PAVs (>100 bp), PAV regions including 2,663 genes based on the criteria that 80% of the region of the gene overlapped the PAV. The majority of PAVs were shorter than 5,000 bp and were notably concentrated within repetitive genomic regions (Figure 3b). In order to elucidate the potential functions of these PAV-associated genes, we conducted an enrichment analysis using the gene annotation results from the eggNOG-mapper tool. We found PAV-associated genes predominately enriched in categories closely related to innate immune responses and immune system processes (Figure S9). These findings suggest that these PAVs may be the one of the reasons for disease resistance in Huizhan in comparison to LTH.

**Figure. 3.**
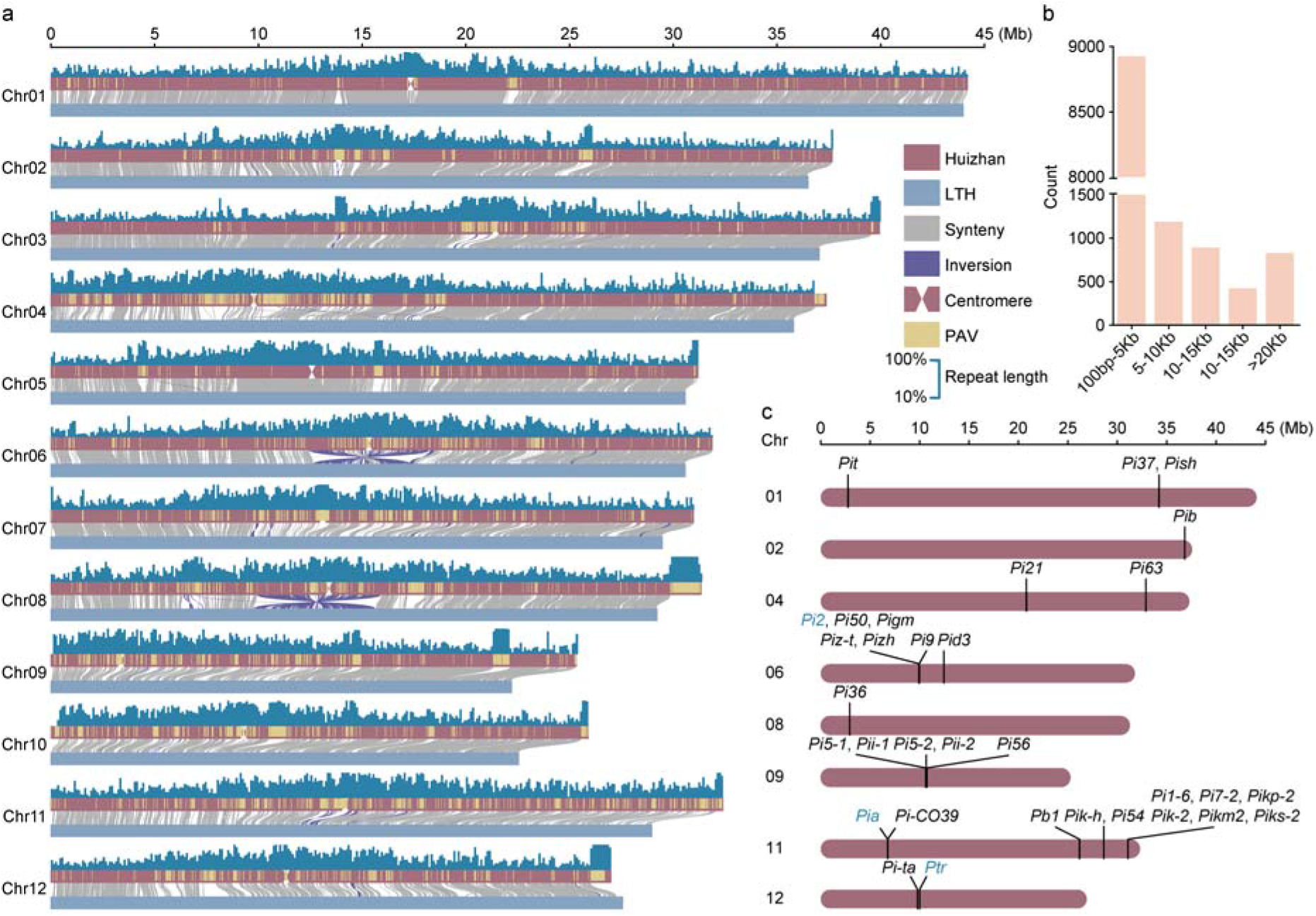
Whole-genome comparison of rice varieties Huizhan and LTH. a. Collinearity analysis of Huizhan and LTH genomes. The collinear regions are depicted as gray lines, and the inversion regions are depicted as purple lines. The presence/absence variations (PAVs) are depicted as yellow blocks. Repeat length percentages are indicated above each chromosome of Huizhan in 100-kb bins. b. Numbers of PAVs of different sizes. c. The genomic position of the *Pi* genes in the Huizhan genome with blue marks for functional *Pi* genes.

### Disease resistance-related genes in Huizhan

Rice blast leads to significant yield losses and adversely impacts rice quality (Sha et al., 2023). NLR proteins, including *Pi* genes, play a key role in activating rice immune responses to rice blast. We systematically categorized *Pi* genes into three main types (Figure 3c, Figure S10, Table S14). Firstly, six *Pi* genes fall into the “absence” category, namely *Pi1-5*, *Pi7-1*, *Pikp-1*, *Pik-1*, *Pikm1*, and *Piks-1*, indicating substantial deficiencies in specific aspects of their genetic composition. Secondly, we identified three *Pi* genes categorized as “functional,” comprising *Pi2*, *Pia* and *Ptr*, indicating their structural integrity (Figure S11, S12 and S13). Lastly, a total of 30 *Pi* genes belong to the “nonfunctional” category. This classification offers profound insights into the diversity and functionality of *Pi* genes within the Huizhan rice variety.

Bacterial blight (BB), caused by *Xanthomonas oryzae* pv. *oryzae* (*Xoo*), is a highly devastating bacterial disease in rice. In a similar approach, we conducted an identification of rice BB resistance genes in the Huizhan. Among the 15 *Xa* genes analysed (Table S14), we successfully identified the functional *Xa27* gene, which remains unaltered at the amino acid level. The *Xa27* gene confers resistance to diverse *Xoo* strains, including the PXO99^A^ strain (Gu et al., 2005). This finding suggests that Huizhan gains part of the resistance to bacterial blight through the *Xa27* gene.

### Comparative analysis of heat stress-responsive transcriptomes

During the heat stress, we observed that Huizhan showed higher fertility than ZH11 (Figure S14a and S14b). Heat stress reduces pollen viability by inhibiting pollen cell growth, abnormal tapetum disintegration and impairment of nutrient storage (Shrestha et al., 2022). To examine whether high temperature affects the pollen development of Huizhan and ZH11, we evaluated their pollen development (Figure 4a). More abnormal pollen grains stained red were observed in ZH11 than in Huizhan. As a result, the pollen vitality of ZH11 was 16.74% lower than that of Huizhan (Figure 4b). We used RNA-seq to investigate the possible underlying factors contributing to heat tolerance of Huizhan. After filtering low-quality reads, we obtained on average 19.30 Gb paired-end reads from each of the three replicates of flower and leaf tissues of Huizhan and ZH11 (Table S15). Using Principal Component Analysis (PCA), we observed that 70.78% of the variance was attributed to the difference between flower and leaf tissues (Figure S15). We performed two comparison groups: Huizhan-flower (HF) vs. ZH11-flower (ZF), Huizhan-leaf (HL) vs. ZH11-leaf (ZL). A total of 6,422 differentially expressed genes (DEGs) were identified in the flower tissue, consisting of 3,367 up-regulated and 3,055 down-regulated genes. Similarly, 6,881 DEGs were detected in the leaf tissue, with 2,940 up-regulated and 3,941 down-regulated genes (Figure 4c). These genes were distributed across all 12 chromosomes of Nipponbare (Supplementary Figure 16). Among them, 1,014 up-regulated genes and 1,416 down-regulated genes were common to both the flower and leaf tissues (Figure 4d). Additionally, Huizhan exhibited fewer down-regulated genes compared to ZH11 in leaf tissues (Figure 4c).

**Figure. 4.**
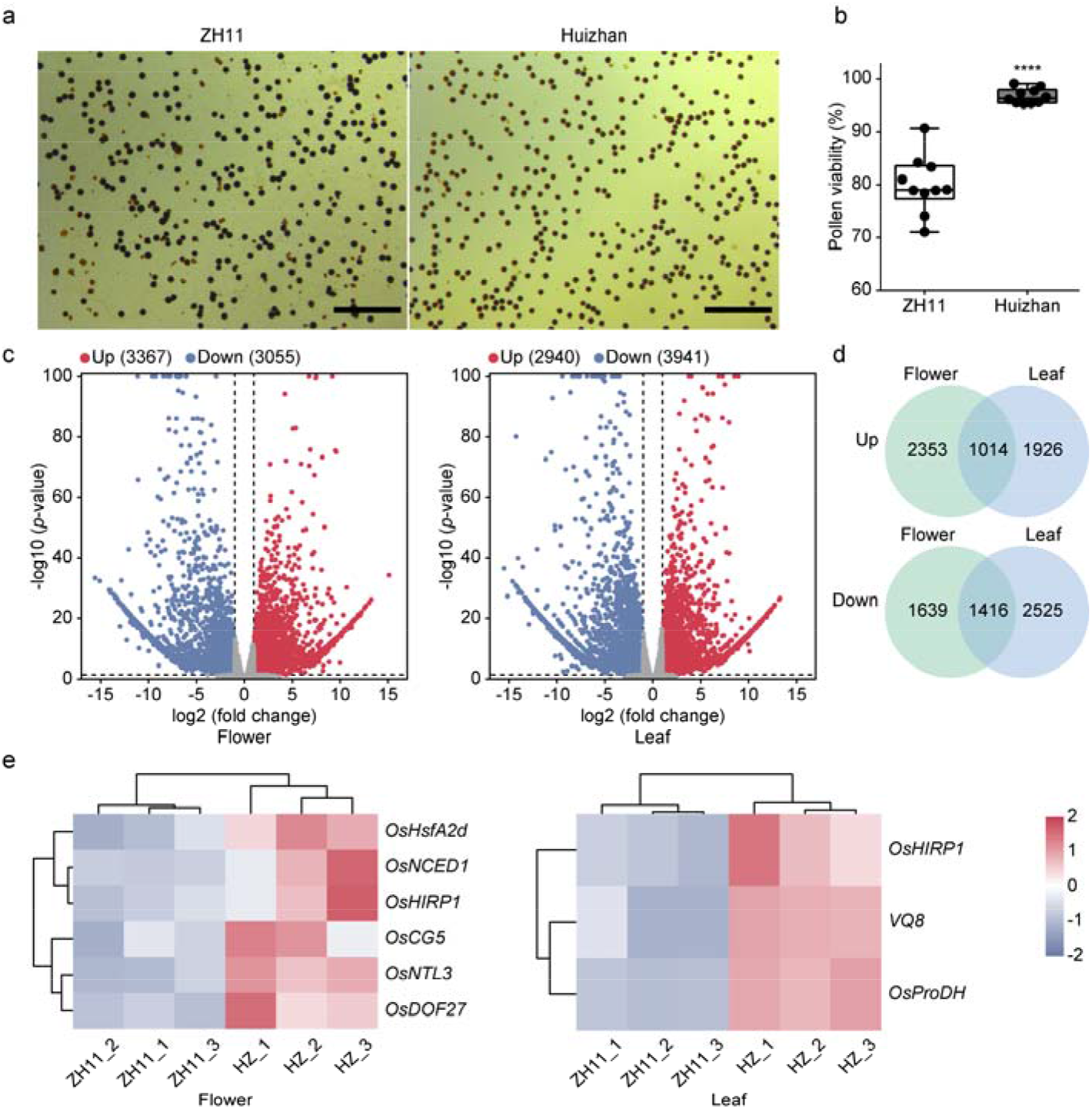
Pollen viability and transcriptomic analysis of heat stress in Huizhan and ZH11. a. Representative pollen staining in Lugol’s solution of ZH11 and Huizhan. Scale bars, 500 μm. b. The comparison of pollen viability of ZH11 and Huizhan. The black solid line in the box represents the median; the upper and lower horizontal lines of the box represent quartiles; the upper and lower adjacent lines represent the minimum and maximum values. Unpaired *t*-test with Welch’s correction was performed. (\*\*\*\**P*L<L0.0001). c. Volcano plots of differentially expressed genes (DEGs) in the flower and leaf tissues under heat stress conditions. d. Venn diagram of up-regulated and down-regulated DEGs shared by flower and leaf tissues. e. The gene expression profiles of genes related to heat tolerance from the funRiceGenes database in rice flower and leaf tissues. The color scale represents the normalized Z-score values of Transcripts Per Million (TPM).

We next performed GO term enrichment analysis of the up-regulated genes. A total of 75 and 20 GO terms were enriched in flower and leaf tissues, respectively (Table S16 and S17). Notably, “response to heat stress” was a commonly enriched GO term in both groups. In the leaf tissue, the enriched GO terms were associated with specific gene functions, including superoxide dismutase activity and positive regulation of transcription in response to heat stress, indicating Huizhan’s active response to mitigate heat stress and maintain cellular homeostasis (Figure S17). Additionally, genes related to responses to biotic stresses were enriched in flower tissues, suggesting Huizhan’s proactive responses to external biological threats such as pathogen infections.

We also performed KEGG pathway enrichment analysis (Bu et al., 2021). In both control groups, up-regulated DEGs were enriched in pathways related to the biosynthesis of secondary metabolites. Recent studies have indicated that pattern-triggered immunity (PTI) and effector-triggered immunity (ETI) may trigger the biosynthesis of a series of similar secondary metabolites and phytohormones that are associated with defense mechanisms (Zhai et al., 2022). This convergence reveals that these pathways may play a regulatory role in the modulation of heat stress response genes in Huizhan (Figure S18). Upon heat treatment, DEGs in flower tissue showed enrichment related to cutin, suberine and wax biosynthesis, highlighting their importance in biological functions. Previous research has shown that cuticular waxes protect plants against a wide range of environmental stresses (Kan et al., 2022). The defective pollen wall from rice (*OsDPW*) (Shi et al., 2011) is associated with the biosynthesis of primary fatty alcohols for the anther cuticle and was found to be significantly up-regulated in the flower of Huizhan. Genes related to heat tolerance were extracted from the funRiceGenes database (Yao et al., 2018). We found that six heat-tolerance genes that were up-regulated in the flower tissue and three in the leaf tissue, respectively. We also found that *OsHIRP1* (Kim et al., 2019) is commonly expressed in both tissue types, indicating its pivotal role in the coordinated response to heat stress (Figure 4e).

### Coordinated regulation of transcription factors coping with heat stress in rice

Rice is highly susceptible to heat stress, particularly during the reproductive stage (Li et al., 2022). To investigate the expression patterns of the heat stress response in Huizhan and ZH11, we focused on representative genes associated with Ca^2+^ signaling, the unfolded protein response (UPR) pathway, regulation of reactive oxygen species (ROS) levels (Superoxide dismutase (SOD) and catalase (CAT)), heat shock factors (HSFs), and other transcription factors (Figure 5).

**Figure. 5.**
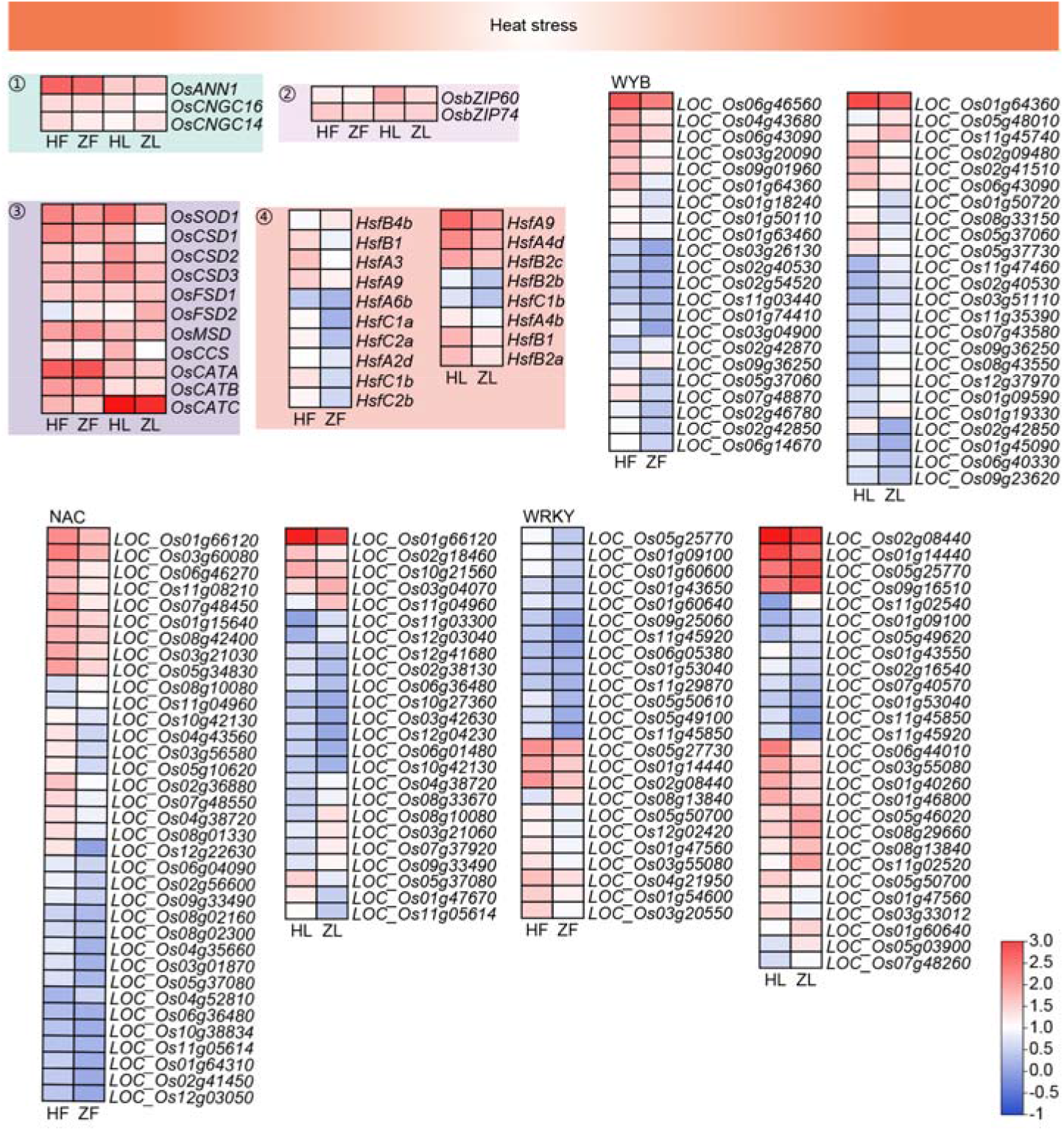
Transcriptome-based analysis of heat tolerance mechanisms in Huizhan and ZH11. Expansion and expression of key genes with the focus on heat-response genes in Huizhan and ZH11. Genes include ①. *OsANN1*, *OsCNGC14* and *OsCNGC16* in the Ca^2+^ signaling; ②. *OsbZIP60* and *OsbZIP74* in the endoplasmic reticulum unfolded-protein response (UPR); ③. Gene family SOD and CAT; ④. Heat shock factors (HSFs). The remaining transcription factors involved in the heat-shock response include NAC, WYB, and WRKY. The heatmap shows the differential gene expression patterns in flower and leaf tissues. The color scale represents the normalized Z-score values of Transcripts Per Million (TPM). HF, Huizhan-flower; ZF, ZH11-flower; HL, Huizhan-leaf; ZL, ZH11-leaf.

Under heat stress, the genes related to Ca^2+^ signaling, including *OsCNGC14*, *OsCNGC16* and *OsANN1*, were expressed in both Huizhan and ZH11. This suggests that both varieties use Ca^2+^ influx as a signaling mechanism in coping with heat stress. Among the UPR pathway genes, *OsbZIP60* was significantly up-regulated in the leaf tissue of Huizhan. Among the CAT and SOD family genes, only *OsFSD2* was down-regulated in flower and leaf tissues, while five SOD genes were up-regulated in leaf tissues of Huizhan. This suggests a more active role of SOD in maintaining ROS homeostasis under heat stress in Huizhan compared to ZH11.

Heat Shock Factors (HSFs) are key transcription factors in cellular stress response and gene expression regulation (Shamshad et al., 2023). In the flower tissue, 10 HSF genes were identified from DEGs, with one down-regulated and nine up-regulated. All eight HSF genes in the leaf tissue exhibited up-regulation (Table S18). Our identified up-regulated HSF genes cover three major branches: A, B and C types (Chauhan et al., 2011). Additionally, transcription factors like NAC, MYB and WRKY are known to enhance heat tolerance in rice (Li et al., 2022). In the flower tissue, we identified 32 NAC genes, 20 WYB genes and 23 WRKY genes as up-regulated genes, and 14 NAC genes, 12 WYB genes and 15 WRKY genes up-regulated in the leaf tissue (Table S19). Notably, the aforementioned nine up-regulated HSF genes in flowers correlated significantly with the majority of NAC genes (18 out of 32), WYB genes (11 out of 20) and nine WRKY genes (*p* < 0.05) in response to heat stress (Table S20). In leaf tissues, three NAC genes, five WYB genes and five WRKY genes showed significant correlation with the eight HSF genes (Table S21). These results suggest that, compared to ZH11, Huizhan relies more on HSFs for improved heat tolerance, and that HSF genes may contribute to the co-regulation of heat tolerance in Huizhan, possibly through their interactions with NAC, WYB and WRKY transcription factors.

## Discussion

In this study, we sought to use genomic and transcriptomic tools to dissect the genetic basis of disease resistance and heat tolerance of the elite rice variety Huizhan. Using Nanopore ultra-long reads and next-generation sequencing short reads, the genome of Huizhan was assembled into 12 pseudomolecules, with the scaffold N50 of 31.87LMb, demonstrating the capacity of the ultra-long reads in generating highly contiguous genome assemblies. We conducted a comprehensive comparative genomic analysis of Huizhan and six other XI rice varieties. Our results identified genomic variations unique to Huizhan. By comparing with LTH, a variety known for its universal susceptibility to rice blast, we identified 12,177 PAVs larger than 100 bp affecting 2,663 genes. The enrichment results of genes overlapping PAVs highlight the potential for improved rice disease resistance in Huizhan. Among the 39 *Pi* genes, three genes, including *Pi2*, *Pia* and *Ptr*, remain complete and unmutated. *Pi2* shows robust resistance against 523 isolates of *M. oryzae* isolated in China (Zhu et al., 2012). *Pia* shows high resistance to rice blast populations in Jiangsu Province, China (Zeng et al., 2011), and serves as a key gene for breeding rice varieties with other rice blast resistance genes (Wu et al., 2015). *Ptr* is another broad-spectrum blast resistance gene (Zhao et al., 2018). Using a similar approach, we identified the functional *Xa27* gene in Huizhan, providing a preliminary explanation for its resistance to bacterial blight. The phylogenetic analysis suggests that Huizhan belongs to a lineage consisting of ZS97 and MH63. Furthermore, we conducted gene family analyses, including the unique gene families. As the shared genes among the GO enrichment results of the non-clustered gene families, OsHZ15485 (*OsARGOS*) (Wang et al., 2009) is related to cell proliferation and cell expansion, suggesting the possible reason for the high-quality yield potential of Huizhan. We also identified gene expansion and contraction. The results of GO enrichment analysis indicate that these genes in Huizhan have an important regulatory function in development and signaling transduction.

Under heat stress conditions, with analyzed DEGs, we conducted GO enrichment analyses and identified genes associated with heat tolerance in Huizhan. KEGG analysis indicated that the increased biosynthesis and accumulation of secondary metabolites in Huizhan may contribute to its enhanced heat tolerance, such as flavonoid playing a role in heat tolerance in rice reproductive tissues (Cai et al., 2020; Shen et al., 2018). Furthermore, Huizhan may alleviate heat stress by increasing the biosynthesis of cutin, suberin and wax in floral organs (Kan et al., 2022), including *OsDPW*, which is involved in anther and pollen development in rice (Shi et al., 2011). Using the funRiceGenes database, we showed the potential role of *OsHIRP1*, highly expressed in flower and leaf tissues, in the response to heat stress. Based on previous research on thermo-perception and thermo-responses in rice, we have observed that both Huizhan and ZH11 respond to high temperature stress through Ca^2+^ signaling. By identifying the SOD gene family, Huizhan was found to have five up-regulated genes in the leaf tissue. Additionally, the *OsbZIP60* encodes a transcriptional activator and induces downstream gene expression in rice plants (Li et al., 2022), is highly expressed in Huizhan.

Transcription factors in plants, particularly HSFs, have a crucial function in the response to heat stress (Guo et al., 2016). Many studies have reported that heat stress tolerance involves not only HSFs but also various other transcription factors including NAC, MYB and WRKY (Li et al., 2022). Our data suggest that Huizhan may respond to heat stress through HSFs. *HsfA2d*, an important regulator of the heat shock response (HSR) in rice (Chauhan et al., 2011), is up-regulated in Huizhan. Similarly, *HsfA9*, induced under heat stress in rice (Mittal et al., 2009), is up-regulated in Huizhan. *HSFA9* from sunflower has been demonstrated to confer heat tolerance in transgenic tobacco plants (Prieto-Dapena et al., 2008). *HsfB1* is another up-regulated gene in Huizhan under heat stress, which has been demonstrated to support the development of acquired thermotolerance during heat stress (Ikeda et al., 2011). Additionally, our study found that Huizhan may coordinate HSFs in the feedback regulation of transcription factors such as NAC, WYB, and WRKY. Thus, there may be a complex interplay and coordination between HSFs and these transcription factors in the regulatory response to heat stress, which could facilitate effective adaptation. However, further investigation is necessary to fully comprehend the mechanisms and implications of this coordination.

In summary, we report the high-quality reference genome of the elite *indica* rice variety Huizhan. The genomic resource is valuable in providing insight into heat adaptation and disease resistance mechanisms in rice. Our study contributes to advancing rice functional genomics studies and facilitates breeding.

## Data availability

Plant Variety Rights (PVR) with Accession No. CNA20191001937 were granted for Huizhan in 2023. Genome and transcriptome sequencing data of Huizhan are available at the National Center for Biotechnology Information under the accession number PRJNA1049505. The genome assembly in this study have been deposited at the Genome Warehouse (GWH) (https://bigd.big.ac.cn/gwh/) through GWHEQVR00000000.The genome annotation file, the chloroplast genome assembly and annotation files are available from Figshare database (https://figshare.com/s/54dc3b710d8683c8e77c).

## Author contributions

G.L., R.H., and W.Y. designed the experiments and their components. R.H. provided all the Huizhan seeds. R.H., Z.Y. and W.Y. collected and cultivated rice Huizhan samples and performed the genome and RNA sequencing. Z.L. examined the pollen viability. G.L. and W.Y. performed genome data analysis, transcriptome analyses, and other data analysis. G.L., W.Y. and L.Y. drafted the manuscript and G.L., R.H., W.Y., L.Y., Z.Z., and T.W. revised the manuscript. All authors read and approved the final manuscript.

## Acknowledgments

We thank the National Key Laboratory of Crop Genetic Improvement at Huazhong Agricultural University for providing the bioinformatics computing platform in this study. This work was supported by STI2030-Major Projects (2023ZD04070), the Key Research & Development Program of Hubei Province (2023BBB171), National Key R&D Program of China (2022YFA1304402), Area Funds of National natural science funds (32160430), Training Program for Academic and Technical Leaders in Major Disciplines of Jiangxi Province (20232BCJ22014), Special Project of Collaborative Innovation in Modern Agricultural Scientific Research of Jiangxi Province (JXXTCX202104), National Natural Science Foundation of China (32172373 and 31801723) and Fundamental Research Funds for the Central Universities (2023ZKPY002, 2662023PY006 and AML2023A05). This work was also supported by Hubei Hongshan Laboratory and National Engineering Research Center of Rice (Nanchang). The authors declare no conflict of interest.

