## Supplementary information for "Genomic and transcriptomic analyses of the elite rice variety Huizhan provide insight into disease resistance and heat tolerance"

**Figure S1. The *k*-mer analysis of the Huizhan genome.**

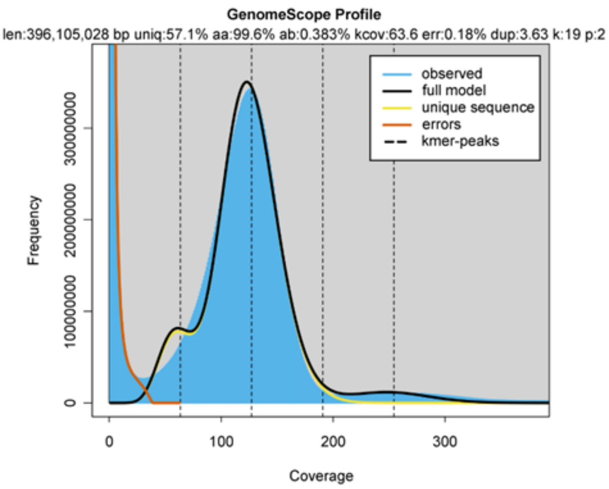


**Figure S2. Distribution of the telomeres and gaps in the Huizhan genome**

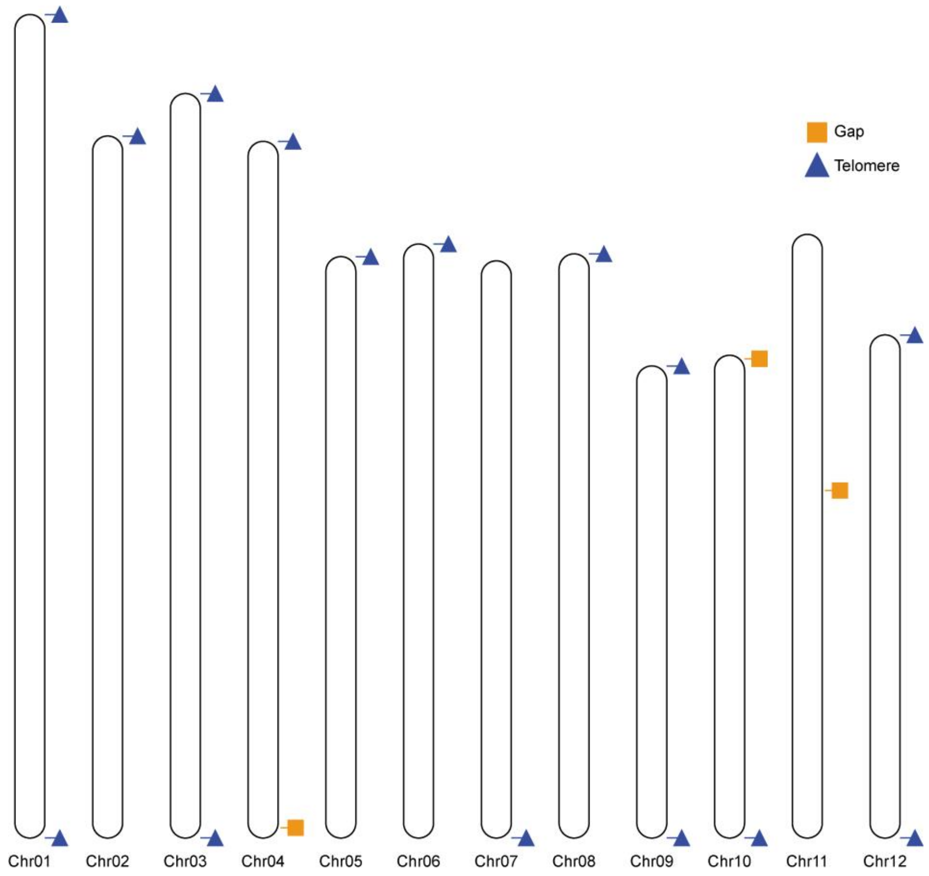


The blue triangles represent the telomeric sequences, while the orange squares indicate the gaps.

**Figure S3. Genome-wide distribution of the ultra-long reads and short reads in the Huizhan genome**

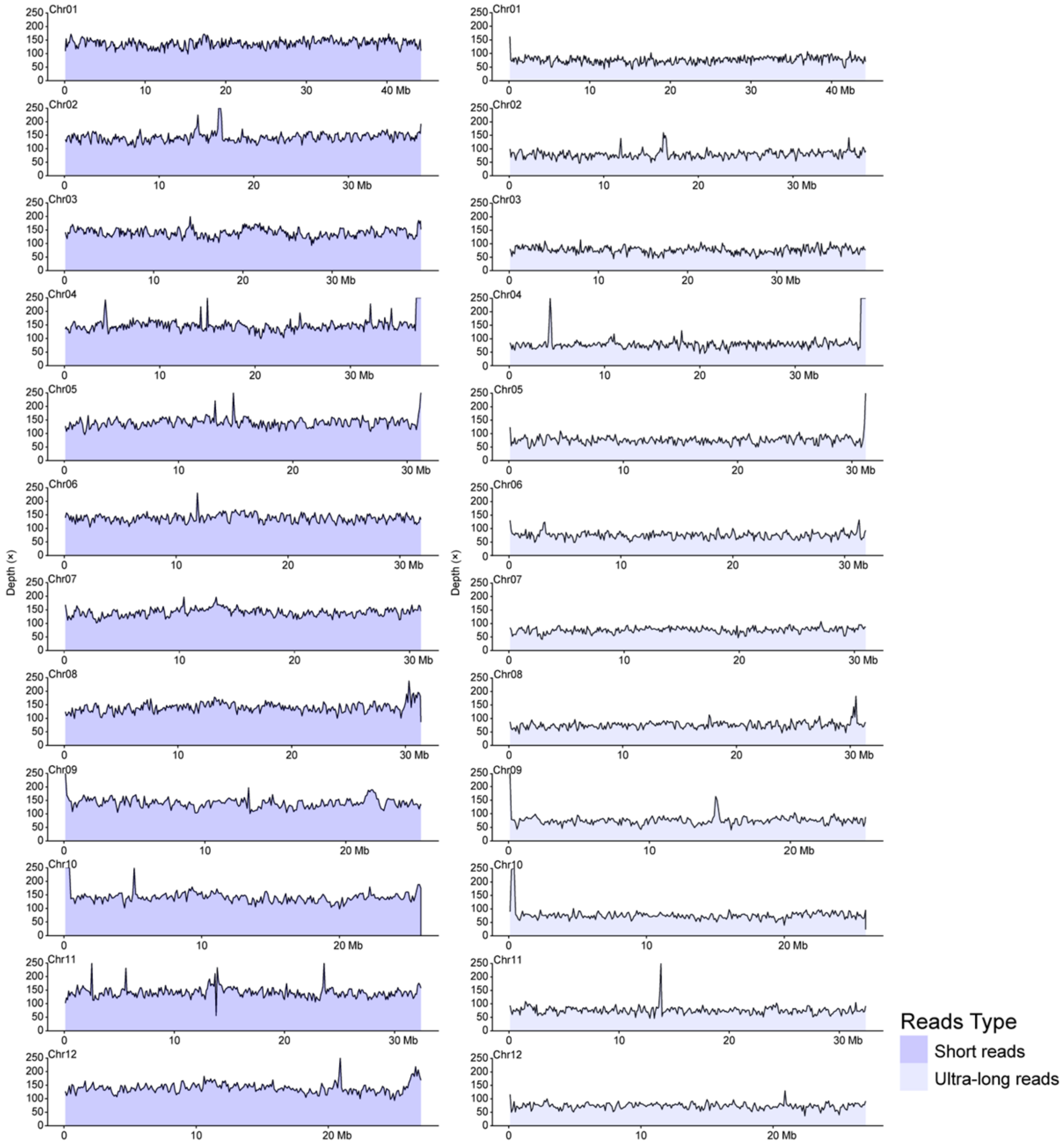


The ultra-long reads were mapped using minimap2 v2.26-r1175, while short reads were mapped using bwa v0.7.17-r1188.

**Figure S4. Map of the Huizhan chloroplast genome**

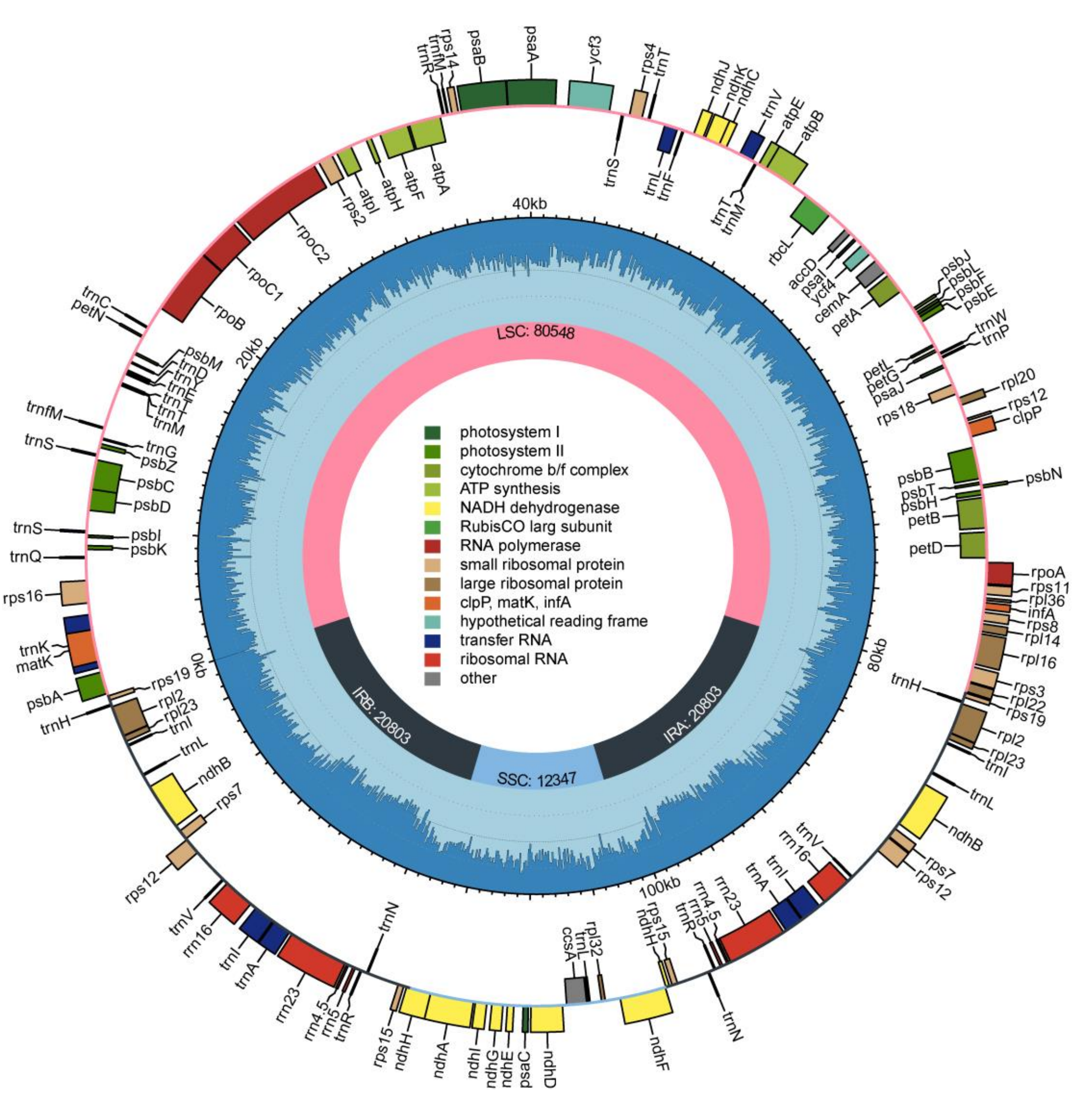


Different colors represent different functions of the genes and the blue inner circle indicates GC content

**Figure S5. Dot plot comparing the Huizhan and six other common XI rice genomes**

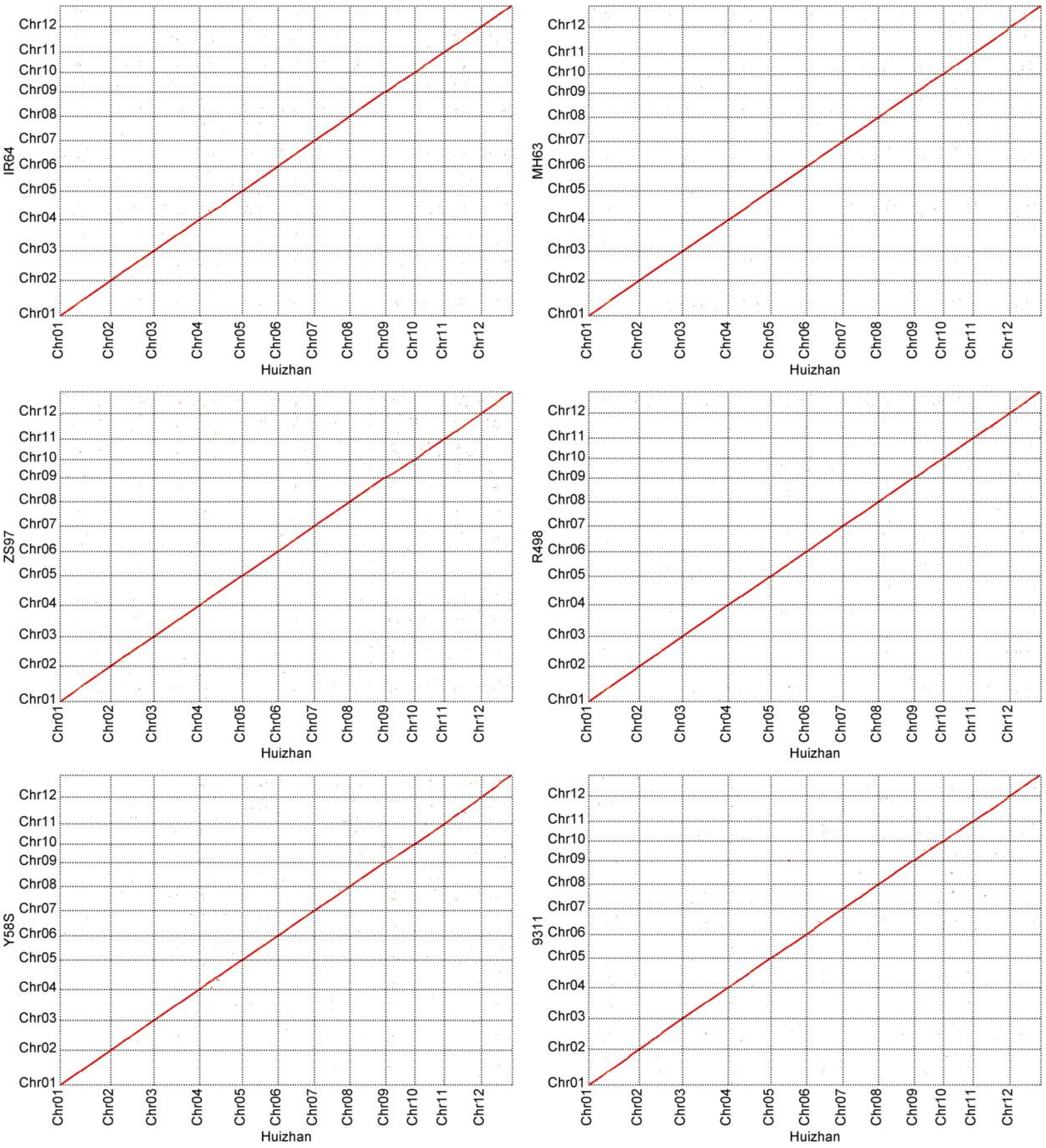

**Figure S6. SNPs and InDels between Huizhan and six other common XI rice genomes**

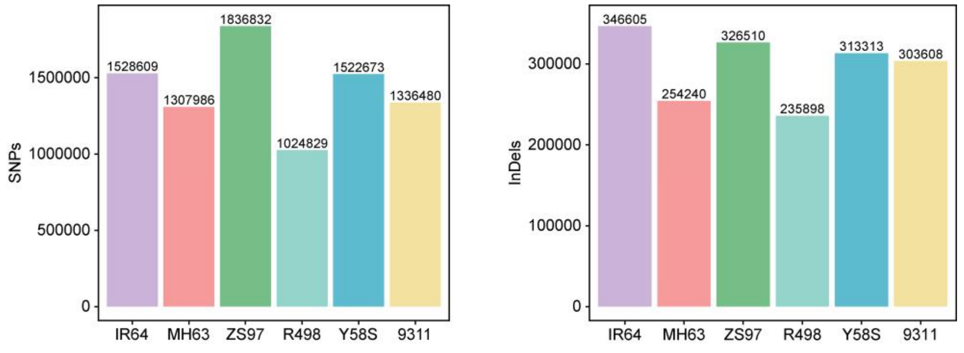


**Figure S7. The density of SNPs and InDels between Huizhan and six other common XI rice genomes**

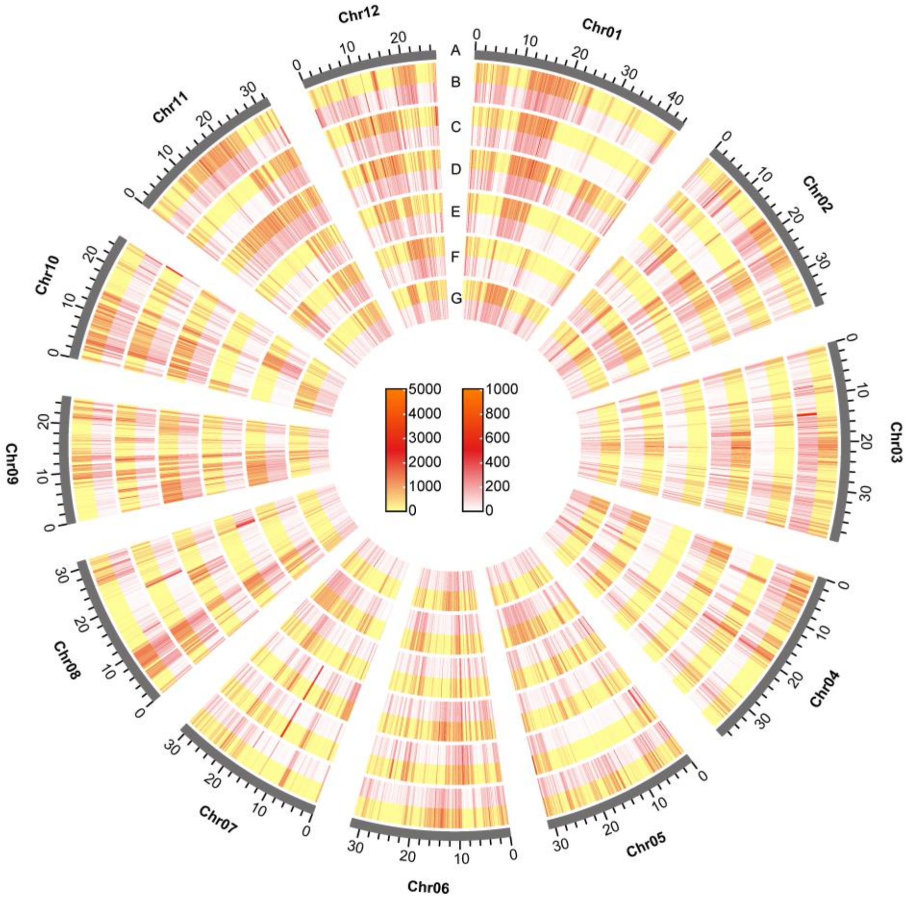


A: The 12 chromosomes, B: IR64, C: MH63, D: ZS97, E: R498, F: Y58S, G: 93-11. Yellow indicates SNPs, while red indicates InDels.

**Figure S8. Collinearity patterns between Huizhan and the corresponding region of six rice accessions of the chromosome 12**

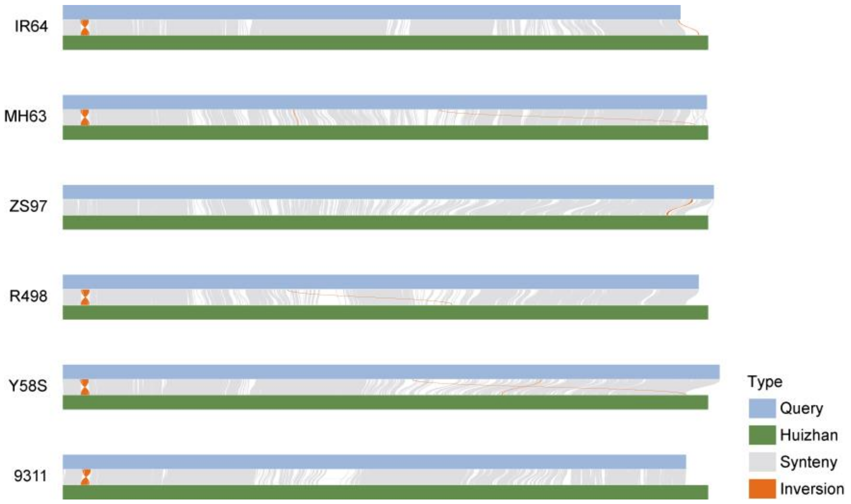


**Figure S9. GO-term enrichment analysis of genes affected by PAVs between Huizhan and LTH**

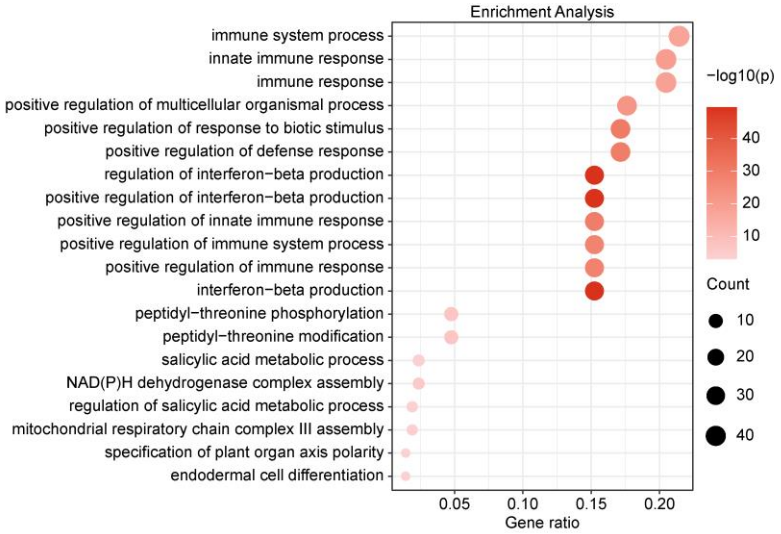

**Figure S10. Comparison of different *Pi* genes in the Huizhan genome.**

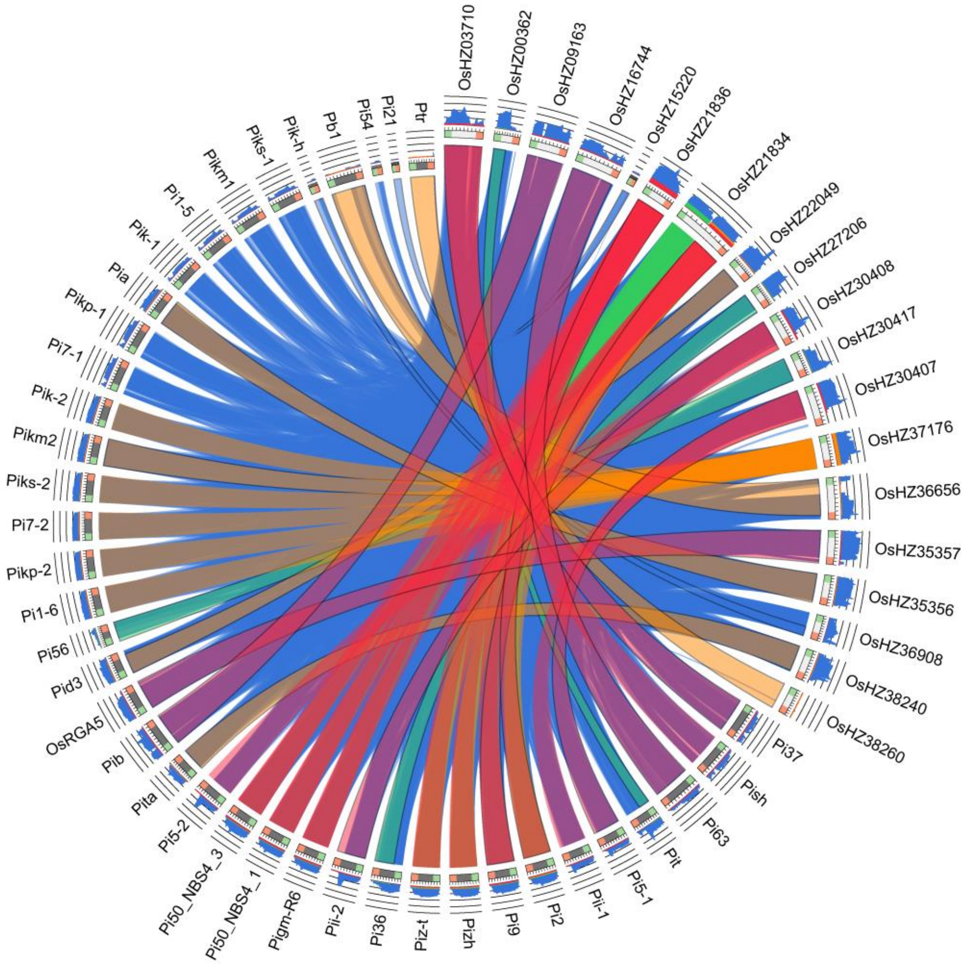


The alignment of Pi genes in LTH genome. The ribbons were colored using the (score-min score) / (max score-min score) ratio with blue<=0.25, green<=0.50, orange<=0.75, red>0.75. The histograms indicate the blast scores.

**Figure S11. Alignment of the candidate Huizhan gene to *Pi2*.**

```
Huizhan_Pi2 (OsHZ21836) MAETVLSMARSLVGSAISKAASAAADETSLLLGVEKDIWYIKDELKTMQAFLRAAELMKK   60
                    Pi2 MAETVLSMARSLVGSAISKAASAAADETSLLLGVEKDIWYIKDELKTMQAFLRAAELMKK   60

Huizhan_Pi2 (OsHZ21836) KDELLKVWAEQIRDLSYDIEDSLDEFKVHIESQTLFRQLVKLRERHRIAIRIHNLKSRVE  120
                    Pi2 KDELLKVWAEQIRDLSYDIEDSLDEFKVHIESQTLFRQLVKLRERHRIAIRIHNLKSRVE  120

Huizhan_Pi2 (OsHZ21836) EVSSRNTRYSLVKPISSGTEIDMDSYAEDIRNQSARNVDEAELVGFSDSKKRLLEMIDTN  180
                    Pi2 EVSSRNTRYSLVKPISSGTEIDMDSYAEDIRNQSARNVDEAELVGFSDSKKRLLEMIDTN  180

Huizhan_Pi2 (OsHZ21836) ANDGPAKVICVVGMGGLGKTALSRKIFESEEDIRKNFPCNAWITVSQSFHRIELLKDMIR  240
                    Pi2 ANDGPAKVICVVGMGGLGKTALSRKIFESEEDIRKNFPCNAWITVSQSFHRIELLKDMIR  240

Huizhan_Pi2 (OsHZ21836) QLLGPSSLDQLLQELQGKVVVQVHHLSEYLIEELKEKRYFVVLDDLWILHDWNWINEIAF  300
                    Pi2 QLLGPSSLDQLLQELQGKVVVQVHHLSEYLIEELKEKRYFVVLDDLWILHDWNWINEIAF  300

Huizhan_Pi2 (OsHZ21836) PKNNKKGSRIVITTRNVDLAEKCATASLVYHLDFLQMNDAITLLLRKTNKNHEDMESNKN  360
                    Pi2 PKNNKKGSRIVITTRNVDLAEKCATASLVYHLDFLQMNDAITLLLRKTNKNHEDMESNKN  360

Huizhan_Pi2 (OsHZ21836) MQKMVERIVNKCGRLPLAILTIGAVLATKQVSEWEKFYEHLPSELEINPSLEALRRMVTL  420
                    Pi2 MQKMVERIVNKCGRLPLAILTIGAVLATKQVSEWEKFYEHLPSELEINPSLEALRRMVTL  420

Huizhan_Pi2 (OsHZ21836) GYNHLPSHLKPCFLYLSIFPEDFEIKRNRLVGRWIAEGFVRPKVGMTTKDVGESYFNELI  480
                    Pi2 GYNHLPSHLKPCFLYLSIFPEDFEIKRNRLVGRWIAEGFVRPKVGMTTKDVGESYFNELI  480

Huizhan_Pi2 (OsHZ21836) NRSMIQRSRVGIAGKIKTCRIHDIIRDITVSISRQENFVLLPMGDGSDLVQENTRHIAFH  540
                    Pi2 NRSMIQRSRVGIAGKIKTCRIHDIIRDITVSISRQENFVLLPMGDGSDLVQENTRHIAFH  540

Huizhan_Pi2 (OsHZ21836) GSMSCKTGLDWSIIRSLAIFGDRPKSLAHAVCPDQLRMLRVLDLEDVTFLITQKDFDRIA  600
                    Pi2 GSMSCKTGLDWSIIRSLAIFGDRPKSLAHAVCPDQLRMLRVLDLEDVTFLITQKDFDRIA  600

Huizhan_Pi2 (OsHZ21836) LLCHLKYLSIGYSSSIYSLPRSIGKLQGLQTLNMPSTYIAALPSEISKLQCLHTLRCIGQ  660
                    Pi2 LLCHLKYLSIGYSSSIYSLPRSIGKLQGLQTLNMPSTYIAALPSEISKLQCLHTLRCIGQ  660

Huizhan_Pi2 (OsHZ21836) FHYDNFSLNHPMKCITNTICLPKVFTPLVSRDDRAKQIAELHMATKSCWSESIGVKVPKG  720
                    Pi2 FHYDNFSLNHPMKCITNTICLPKVFTPLVSRDDRAKQIAELHMATKSCWSESIGVKVPKG  720

Huizhan_Pi2 (OsHZ21836) IGKLRDLQVLEYVDIRRTSSRAIKELGQLSKLRKLGVTTNGSTKEKCKILYAAIEKLSSL  780
                    Pi2 IGKLRDLQVLEYVDIRRTSSRAIKELGQLSKLRKLGVTTNGSTKEKCKILYAAIEKLSSL  780

Huizhan_Pi2 (OsHZ21836) QSLHVDAAGISDGGTLECLDSISSPPPLLRTLVLDGILEEMPNWIEQLTHLKKIYLLRSK  840
                    Pi2 QSLHVDAAGISDGGTLECLDSISSPPPLLRTLVLDGILEEMPNWIEQLTHLKKIYLLRSK  840

Huizhan_Pi2 (OsHZ21836) LKEGKTMLILGALPNLMVLHLYRNAYLGEKLVFKTGAFPNLRTLWIYELDQLREIRFEDG  900
                    Pi2 LKEGKTMLILGALPNLMVLHLYRNAYLGEKLVFKTGAFPNLRTLWIYELDQLREIRFEDG  900

Huizhan_Pi2 (OsHZ21836) SSPLLEKIEIGECRLESGITGIIHLPKLKEIPIRYGSKVAGLGQLEGEVNAHPNRPVLLM  960
                    Pi2 SSPLLEKIEIGECRLESGITGIIHLPKLKEIPIRYGSKVAGLGQLEGEVNAHPNRPVLLM  960

Huizhan_Pi2 (OsHZ21836) YSDRRYHDLGAEAEGSSIEVQTADPVPDAEGSVTVAVEATDPLPEQEGESSQSQVITLTT 1020
                    Pi2 YSDRRYHDLGAEAEGSSIEVQTADPVPDAEGSVTVAVEATDPLPEQEGESSQSQVITLTT 1020

Huizhan_Pi2 (OsHZ21836) NDSEEIGTAQAG                                                 1032
                    Pi2 NDSEEIGTAQAG                                                 1032
```

**Figure S12. Alignment of the candidate Huizhan gene to *Pib***

```
Huizhan_Pib (OsHZ09163) MEATALSVGKSVLNGALGYAKSAFAEEVALQLGIQKDHTFVADELEMMRSFMMEAHEEQD   60
                    Pib MEATALSVGKSVLNGALGYAKSAFAEEVALQLGIQKDHTFVADELEMMRSFMMEAHEEQD   60

Huizhan_Pib (OsHZ09163) NSKVVKTWVKQVRDTAYDVEDSLQDFAVHLKRPSWWRFPRTLLERHRVAKQMKELRNKVE  120
                    Pib NSKVVKTWVKQVRDTAYDVEDSLQDFAVHLKRPSWWRFPRTLLERHRVAKQMKELRNKVE  120

Huizhan_Pib (OsHZ09163) DVSQRNVRYHLIKGSAKATINSTEQSSVIATAIFGIDDARRAAKQDNQRVDLVQLINSED  180
                    Pib DVSQRNVRYHLIKGSAKATINSTEQSSVIATAIFGIDDARRAAKQDNQRVDLVQLINSED  180

Huizhan_Pib (OsHZ09163) QDLKVIAVWGTSGDMGQTTIIRMAYENPDVQIRFPCRAWVRVMHPFSPRDFVQSLVNQLH  240
                    Pib QDLKVIAVWGTSGDMGQTTIIRMAYENPDVQIRFPCRAWVRVMHPFSPRDFVQSLVNQLH  240

Huizhan_Pib (OsHZ09163) ATQGVEALLEKEKTEQDLAKKFNGCVNDRKCLIVLNDLSTIEEWDQIKKCFQKCRKGSRI  300
                    Pib ATQGVEALLEKEKTEQDLAKKFNGCVNDRKCLIVLNDLSTIEEWDQIKKCFQKCRKGSRI  300

Huizhan_Pib (OsHZ09163) IVSSTQVEVASLCAGQESQASELKQLSADQTLYAFYDKGSQIIEDSVKPVSISDVAITST  360
                    Pib IVSSTQVEVASLCAGQESQASELKQLSADQTLYAFYDKGSQIIEDSVKPVSISDVAITST  360

Huizhan_Pib (OsHZ09163) NNHTVAHGEIIDDQSMDADEKKVARKSLTRIRTSVGASEESQLIGREKEISEITHLILNN  420
                    Pib NNHTVAHGEIIDDQSMDADEKKVARKSLTRIRTSVGASEESQLIGREKEISEITHLILNN  420

Huizhan_Pib (OsHZ09163) DSQQVQVISVWGMGGLGKTTLVSGVYQSPRLSDKFDKYVFVTIMRPFILVELLRSLAEQL  480
                    Pib DSQQVQVISVWGMGGLGKTTLVSGVYQSPRLSDKFDKYVFVTIMRPFILVELLRSLAEQL  480

Huizhan_Pib (OsHZ09163) HKGSSKKEELLENRVSSKKSLASMEDTELTGQLKRLLEKKSCLIVLDDFSDTSEWDQIKP  540
                    Pib HKGSSKKEELLENRVSSKKSLASMEDTELTGQLKRLLEKKSCLIVLDDFSDTSEWDQIKP  540

Huizhan_Pib (OsHZ09163) TLFPLLEKTSRIIVTTRKENIANHCSGKNGNVHNLKVLKHNDALCLLSEKVFEEATYLDD  600
                    Pib TLFPLLEKTSRIIVTTRKENIANHCSGKNGNVHNLKVLKHNDALCLLSEKVFEEATYLDD  600

Huizhan_Pib (OsHZ09163) QNNPELVKEAKQILKKCDGLPLAIVVIGGFLANRPKTPEEWRKLNENINAELEMNPELGM  660
                    Pib QNNPELVKEAKQILKKCDGLPLAIVVIGGFLANRPKTPEEWRKLNENINAELEMNPELGM  660

Huizhan_Pib (OsHZ09163) IRTVLEKSYDGLPYHLKSCFLYLSIFPEDQIISRRRLVHRWAAEGYSTAAHGKSAIEIAN  720
                    Pib IRTVLEKSYDGLPYHLKSCFLYLSIFPEDQIISRRRLVHRWAAEGYSTAAHGKSAIEIAN  720

Huizhan_Pib (OsHZ09163) GYFMELKNRSMILPFQQSGSSRKSIDSCKVHDLMRDIAISKSTEENLVFRVEEGCSAYIH  780
                    Pib GYFMELKNRSMILPFQQSGSSRKSIDSCKVHDLMRDIAISKSTEENLVFRVEEGCSAYIH  780

Huizhan_Pib (OsHZ09163) GAIRHLAISSNWKGDKSEFEGIVDLSRIRSLSLFGDWKPFFVYGKMRFIRVLDFEGTRGL  840
                    Pib GAIRHLAISSNWKGDKSEFEGIVDLSRIRSLSLFGDWKPFFVYGKMRFIRVLDFEGTRGL  840

Huizhan_Pib (OsHZ09163) EYHHLDQIWKLNHLKFLSLRGCYRIDLLPDLLGNLRQLQMLDIRGTYVKALPKTIIKLQK  900
                    Pib EYHHLDQIWKLNHLKFLSLRGCYRIDLLPDLLGNLRQLQMLDIRGTYVKALPKTIIKLQK  900

Huizhan_Pib (OsHZ09163) LQYIHAGRKTDYVWEEKHSLMQRCRKVGCICATCCLPLLCEMYGPLHKALARRDAWTFAC  960
                    Pib LQYIHAGRKTDYVWEEKHSLMQRCRKVGCICATCCLPLLCEMYGPLHKALARRDAWTFAC  960

Huizhan_Pib (OsHZ09163) CVKFPSIMTGVHEEEGAMVPSGIRKLKDLHTLRNINVGRGNAILRDIGMLTGLHKLGVAG 1020
                    Pib CVKFPSIMTGVHEEEGAMVPSGIRKLKDLHTLRNINVGRGNAILRDIGMLTGLHKLGVAG 1020

Huizhan_Pib (OsHZ09163) INKKNGRAFRLAISNLNKLESLSVSSAGMPGLCGCLDDISSPPENLQSLKLYGSLKTLPE 1080
                    Pib INKKNGRAFRLAISNLNKLESLSVSSAGMPGLCGCLDDISSPPENLQSLKLYGSLKTLPE 1080

Huizhan_Pib (OsHZ09163) WIKELQHLVKLKLVSTRLLEHDVAMEFLGELPKVEILVISPFKSEEIHFKPPQTGTAFVS 1140
                    Pib WIKELQHLVKLKLVSTRLLEHDVAMEFLGELPKVEILVISPFKSEEIHFKPPQTGTAFVS 1140

Huizhan_Pib (OsHZ09163) LRVLKLAGLWGIKSVKFEEGTMPKLERLQVQGRIENEIGFSGLEFLQNINEVQLSVWFPT 1200
                    Pib LRVLKLAGLWGIKSVKFEEGTMPKLERLQVQGRIENEIGFSGLEFLQNINEVQLSVWFPT 1200

Huizhan_Pib (OsHZ09163) DHDRIRAARAAGADYETAWEEEVQEARRKGGELKRKIREQLARNPNQPIIT          1251
                    Pib DHDRIRAARAAGADYETAWEEEVQEARRKGGELKRKIREQLARNPNQPIIT          1251
```

**Figure S13. Alignment of the candidate Huizhan gene to *Ptr***

```
Huizhan_Ptr (OsHZ38260) MDRLWAAPPLSSPLSYPVARGGGAWPTHPAAELGCLAEEGSMSGAGQSRGHRLGLHIDSD  60
                    Ptr MDRLWAAPPLSSPLSYPVARGGGAWPTHPAAELGCLAEEGSMSGAGQSRGHRLGLHIDSD  60

Huizhan_Ptr (OsHZ38260) WPEVLLINDYAVFMGYLSMVVTGTGFLVLTWSTVILLGGFVSMLSNKDFWSLTVITLVQT 120
                    Ptr WPEVLLINDYAVFMGYLSMVVTGTGFLVLTWSTVILLGGFVSMLSNKDFWSLTVITLVQT 120

Huizhan_Ptr (OsHZ38260) RIFDVFLNGKVSHIGYSLKRLCKAARFIALPHNHKKVGFRGAVRVLVFTIVLCPLFLLYM 180
                    Ptr RIFDVFLNGKVSHIGYSLKRLCKAARFIALPHNHKKVGFRGAVRVLVFTIVLCPLFLLYM 180

Huizhan_Ptr (OsHZ38260) FGLFVSPWISLWRLIQQDYGVTAGDSSSKAHLQPALVVLYSLALFQGVLFYYRAISAWEE 240
                    Ptr FGLFVSPWISLWRLIQQDYGVTAGDSSSKAHLQPALVVLYSLALFQGVLFYYRAISAWEE 240

Huizhan_Ptr (OsHZ38260) QKLVKDVADKYMFDTVSRSSVSDYLHEIKVGCENDPSFARGRNLITYAVKLMESTSPDGY 300
                    Ptr QKLVKDVADKYMFDTVSRSSVSDYLHEIKVGCENDPSFARGRNLITYAVKLMESTSPDGY 300

Huizhan_Ptr (OsHZ38260) LSGARILDTLIKFNRDDASGSELPGQSMQIYNMIGSASSSPILHNLVQMLDFKSAYDGEI 360
                    Ptr LSGARILDTLIKFNRDDASGSELPGQSMQIYNMIGSASSSPILHNLVQMLDFKSAYDGEI 360

Huizhan_Ptr (OsHZ38260) RLRAARIVEHFAGEVRLDKILQGIRCVSSLLELEQKGFQNDHHSSFQEDDGDQLSFEEED 420
                    Ptr RLRAARIVEHFAGEVRLDKILQGIRCVSSLLELEQKGFQNDHHSSFQEDDGDQLSFEEED 420

Huizhan_Ptr (OsHZ38260) DHQISVKEKDYYPKDYKQMQLTGMQILLKLSYDKNNLFLMSNTDDPALINKIVALITSKG 480
                    Ptr DHQISVKEKDYYPKDYKQMQLTGMQILLKLSYDKNNLFLMSNTDDPALINKIVALITSKG 480

Huizhan_Ptr (OsHZ38260) SLHKKQHNEWSCMAELGVKILSRFMRFMYGPTKSNNILWHEISTSSKAIGTLESILECDQ 540
                    Ptr SLHKKQHNEWSCMAELGVKILSRFMRFMYGPTKSNNILWHEISTSSKAIGTLESILECDQ 540

Huizhan_Ptr (OsHZ38260) CDSVLKKHAIRILKRIFMDTSSAMGEGDRERFIGSLMDMSLHNSNGDFQNLAGVDLALKK 600
                    Ptr CDSVLKKHAIRILKRIFMDTSSAMGEGDRERFIGSLMDMSLHNSNGDFQNLAGVDLALKK 600

Huizhan_Ptr (OsHZ38260) QGLSILKEIYLNPSSIMGEGDRERFIGSLMDMFLDNSKGDFGNLPGEDLDLKKQELSILK 660
                    Ptr QGLSILKEIYLNPSSIMGEGDRERFIGSLMDMFLDNSKGDFGNLPGEDLDLKKQELSILK 660

Huizhan_Ptr (OsHZ38260) EICMDPSSFMGEGDREKFIGTLMDMFLHNSKGDLFEKLAGDDLVQICRRSGSSATIILRK 720
                    Ptr EICMDPSSFMGEGDREKFIGTLMDMFLHNSKGDLFEKLAGDDLVQICRRSGSSATIILRK 720

Huizhan_Ptr (OsHZ38260) YGHDIVDCIADTRSSVYSSMHRKIAAKILNHLCSPYSTDEEHLQNLKEAIIDLIPKVLRE 780
                    Ptr YGHDIVDCIADTRSSVYSSMHRKIAAKILNHLCSPYSTDEEHLQNLKEAIIDLIPKVLRE 780

Huizhan_Ptr (OsHZ38260) ALGWGLTGEEILRVAVSGLEGTQDDDWKLQEALASLCATVFNRIVSKDADLTARFNNIAA 840
                    Ptr ALGWGLTGEEILRVAVSGLEGTQDDDWKLQEALASLCATVFNRIVSKDADLTARFNNIAA 840

Huizhan_Ptr (OsHZ38260) GICDQTTKPRMTFADLINEAVKVHRIEFMPEPPAKPEPYEFMPAKYPPPHYMFLVEEDPN 900
                    Ptr GICDQTTKPRMTFADLINEAVKVHRIEFMPEPPAKPEPYEFMPAKYPPPHYMFLVEEDPN 900

Huizhan_Ptr (OsHZ38260) ACCIS                                                        905
                    Ptr ACCIS                                                        905
```

**Figure S14. Phenotypes of Huizhan and ZH11 in response to heat stress treatments**

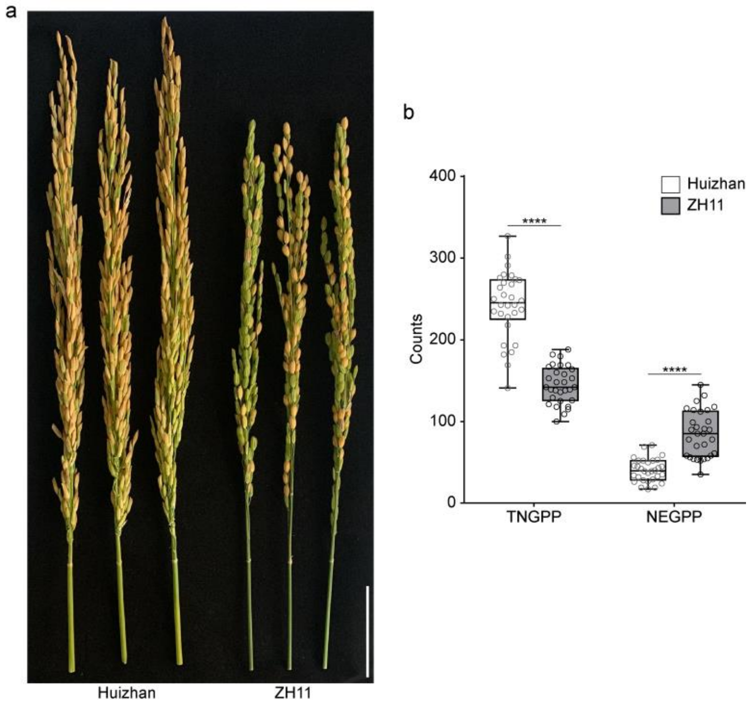


a. The panicle of Huizhan and ZH11 under heat stress. Scale bar, 5cm. b. The evaluation of fertility ability. TNGPP, total number of grains per panicle. NEGPP, number of empty grains per panicle. Asterisks indicate significant differences using the two-tailed Student's *t*-test (*****P* < 0.0001). The box plot elements are: centre line, median; box limits, 25th and 75th percentiles.

**Figure S15. Principal component analysis (PCA) analysis of the RNA-seq data of Huizhan and ZH11**

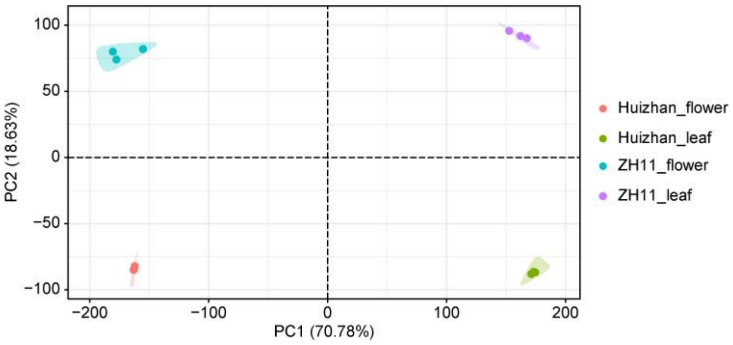

**Figure S16. Genome-wide distribution of Differentially Expressed Genes (DEGs) in the reference genome Nipponbare.**

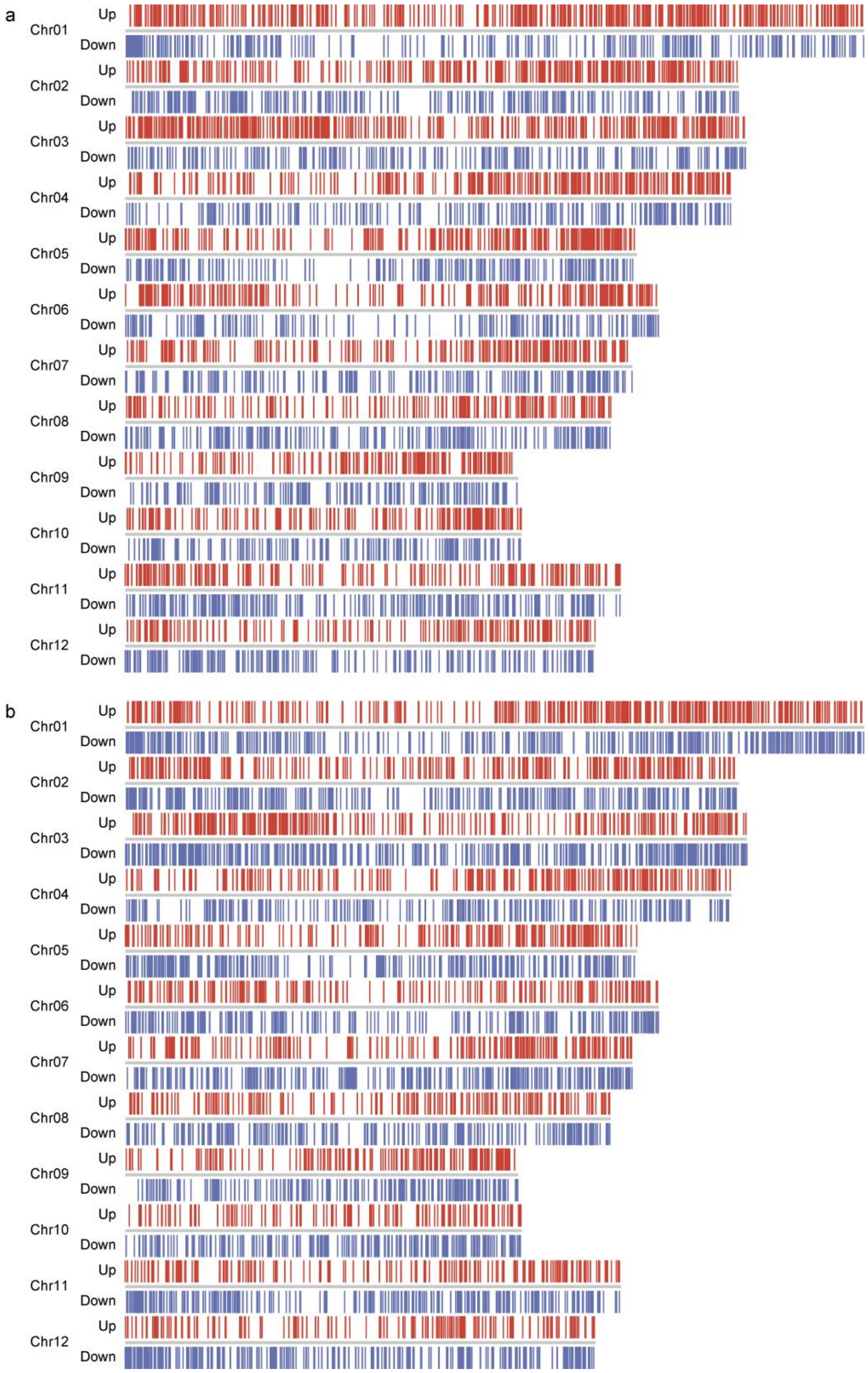


a and b represent DEGs in flower and leaf, respectively. Up-regulated genes are shown in red, while down-regulated genes are shown in blue.

**Figure S17. Go-term enrichment analysis of up-regulated genes**

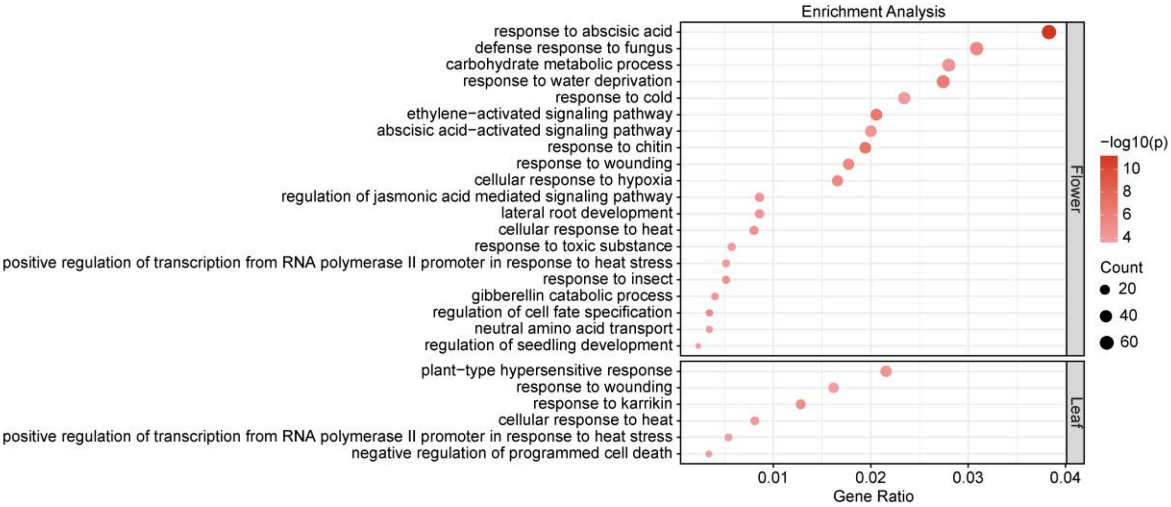


**Figure S18. KEGG pathway enrichment analysis of up-regulated genes**

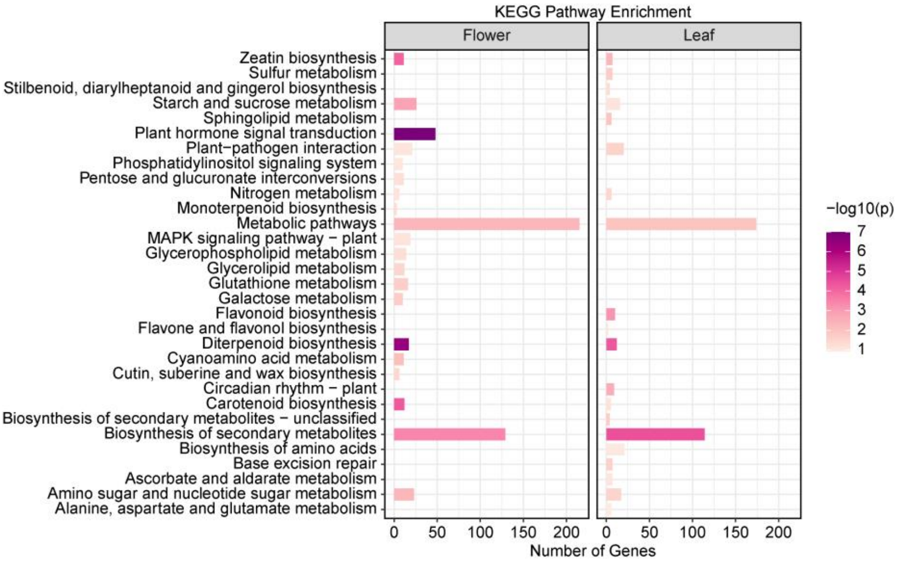

**Table S1. Analysis of the Ultra-long reads and short sequencing reads used for genome assembly**

| Types | Platform | Clean reads | Clean base (Gb) | Depth (X) |
|---|---|---|---|---|
| Ultra-long reads | Promethion | 808,518 | 31.6 | 91 |
| Short reads | DNBSEQ-T7 | 396,246,064 | 59.4 | 149 |

**Table S2. Analysis of the RNA-seq data used for genome annotation**

| Tissue | Platform | Clean reads | Clean base (Gb) |
|---|---|---|---|
| Roots | DNBSEQ-T7 | 46,288,566 | 6.9 |
| Leaves | DNBSEQ-T7 | 77,597,820 | 11.6 |
| Flag leaf | DNBSEQ-T7 | 64,059,126 | 9.6 |
| Shoots | DNBSEQ-T7 | 80,598,842 | 12.1 |
| Panicles | DNBSEQ-T7 | 64,127,892 | 9.6 |
| Ear (before heading) | DNBSEQ-T7 | 63,245,982 | 9.5 |
| Ear (after heading) | DNBSEQ-T7 | 69,540,518 | 10.4 |

**Table S3. Statistics of the chromosome size of the Huizhan genome**

| Chromosome | Length | GC |
|---|---|---|
| Chr01 | 44,187,703 | 43.66 |
| Chr02 | 37,668,565 | 43.42 |
| Chr03 | 39,948,252 | 43.87 |
| Chr04 | 37,381,897 | 43.97 |
| Chr05 | 31,196,619 | 43.96 |
| Chr06 | 31,873,801 | 43.47 |
| Chr07 | 30,971,594 | 43.51 |
| Chr08 | 31,345,896 | 43.69 |
| Chr09 | 25,330,080 | 43.92 |
| Chr10 | 25,904,695 | 43.82 |
| Chr11 | 32,390,624 | 43.42 |
| Chr12 | 26,996,961 | 43.04 |

**Table S4. The *CentO*-enriched regions of the Huizhan genome**

| Chromosome | Start | End |
|---|---|---|
| Chr01 | 16,948,210 | 17,735,704 |
| Chr02 | 13,610,356 | 14,140,459 |
| Chr03 | 21,214,785 | 21,638,655 |
| Chr04 | 9,699,432 | 9,858,988 |
| Chr05 | 12,448,365 | 12,707,441 |
| Chr06 | 15,219,460 | 15,421,242 |
| Chr07 | 12,716,022 | 13,487,494 |
| Chr08 | 13,140,846 | 13,661,306 |
| Chr09 | 3,115,237 | 3,640,510 |
| Chr10 | 8,955,114 | 9,622,738 |
| Chr11 | 13,958,925 | 14,406,688 |
| Chr12 | 11,223,477 | 11,468,311 |

**Table S5. Statistics of the Telomere region in the Huizhan genome.**

| Chromosome | Status | Left telomere number | Right telomere number |
|---|---|---|---|
| Chr01 | both | 133 | 1,716 |
| Chr02 | left | 1,314 | 0 |
| Chr03 | both | 1,195 | 1,055 |
| Chr04 | left | 252 | 0 |
| Chr05 | left | 1,901 | 0 |
| Chr06 | left | 1,926 | 0 |
| Chr07 | right | 0 | 1,585 |
| Chr08 | left | 954 | 0 |
| Chr09 | both | 1,051 | 664 |
| Chr10 | right | 0 | 897 |
| Chr11 | no | 0 | 0 |
| Chr12 | both | 152 | 1,165 |

**Table S6. BUSCO assessment of the Huizhan genome**

| Type | Huizhan-assembly | Huizhan-annotation |
|---|---|---|
| Complete BUSCOs (C) | 1,592 (98.7%) | 1,573 (97.4%) |
| Complete and single-copy BUSCOs (S) | 1,557 (96.5%) | 1,537 (95.2%) |
| Complete and duplicated BUSCOs (D) | 35 (2.2%) | 36 (2.2%) |
| Fragmented BUSCOs (F) | 13 (0.8%) | 13 (0.8%) |
| Missing BUSCOs (M) | 9 (0.5%) | 28 (1.8%) |
| Total BUSCO groups searched | 1614 | 1614 |

**Table S7. Consensus quality value (QV) score of the Huizhan genome**

| Chromosome | k_asm | k_total | Error rate | QV |
|---|---|---|---|---|
| Chr01 | 86,895 | 44,187,685 | 0.000103596 | 39.8466 |
| Chr02 | 63,384 | 37,668,547 | 8.86326E-05 | 40.5241 |
| Chr03 | 62,276 | 39,948,234 | 8.21089E-05 | 40.8561 |
| Chr04 | 91,597 | 37,381,761 | 0.000129114 | 38.8903 |
| Chr05 | 53,892 | 31,196,601 | 9.09953E-05 | 40.4098 |
| Chr06 | 69,254 | 31,873,783 | 0.000114474 | 39.4129 |
| Chr07 | 57,212 | 30,971,576 | 9.73084E-05 | 40.1185 |
| Chr08 | 101,199 | 31,345,878 | 0.000170179 | 37.6909 |
| Chr09 | 48,500 | 25,330,062 | 0.000100866 | 39.9625 |
| Chr10 | 43,138 | 25,904,559 | 8.77148E-05 | 40.5693 |
| Chr11 | 111,073 | 32,390,488 | 0.000180777 | 37.4286 |
| Chr12 | 51,941 | 26,996,943 | 0.000101353 | 39.9416 |
| Whole genome | 840,361 | 395,196,117 | 0.000112031 | 39.5066 |

**Table S8. Codon used in the chloroplast genome assembly of Huizhan**

| Codon | AA | Fraction | Frequency | Number |
|---|---|---|---|---|
| AAA | K | 0.641 | 40.796 | 1,829 |
| TTT | F | 0.579 | 39.19 | 1,757 |
| AAT | N | 0.649 | 30.58 | 1,371 |
| ATT | I | 0.406 | 29.376 | 1,317 |
| TTC | F | 0.421 | 28.506 | 1,278 |
| GAA | E | 0.666 | 27.19 | 1,219 |
| TAT | Y | 0.634 | 25.495 | 1,143 |
| ATA | I | 0.337 | 24.335 | 1,091 |
| AAG | K | 0.359 | 22.863 | 1,025 |
| AGA | R | 0.326 | 22.283 | 999 |
| CTT | L | 0.211 | 21.837 | 979 |
| TCT | S | 0.238 | 21.547 | 966 |
| TAA | * | 0.385 | 21.524 | 965 |
| TTA | L | 0.205 | 21.19 | 950 |
| CAA | Q | 0.689 | 20.654 | 926 |
| TTG | L | 0.197 | 20.431 | 916 |
| ATC | I | 0.257 | 18.58 | 833 |
| GAT | D | 0.667 | 17.978 | 806 |
| TCC | S | 0.194 | 17.576 | 788 |
| TAG | * | 0.312 | 17.465 | 783 |
| CAT | H | 0.681 | 17.019 | 763 |
| TGA | * | 0.302 | 16.907 | 758 |
| GGA | G | 0.347 | 16.573 | 743 |
| CTA | L | 0.16 | 16.573 | 743 |
| AAC | N | 0.351 | 16.528 | 741 |
| TCA | S | 0.179 | 16.149 | 724 |
| TGG | W | 1 | 15.435 | 692 |
| ATG | M | 1 | 15.368 | 689 |
| GTT | V | 0.331 | 15.212 | 682 |
| TAC | Y | 0.366 | 14.744 | 661 |
| AGG | R | 0.215 | 14.677 | 658 |
| CCA | P | 0.308 | 14.632 | 656 |
| GTA | V | 0.31 | 14.231 | 638 |
| CTC | L | 0.133 | 13.784 | 618 |
| GAG | E | 0.334 | 13.628 | 611 |
| AGT | S | 0.149 | 13.495 | 605 |
| GGG | G | 0.279 | 13.316 | 597 |
| CCT | P | 0.278 | 13.205 | 592 |
| ACT | T | 0.305 | 12.848 | 576 |
| TGT | C | 0.551 | 12.134 | 544 |

| | | | | |
|---|---|---|---|---|
| CCC | P | 0.25 | 11.889 | 533 |
| CGA | R | 0.166 | 11.353 | 509 |
| ACC | T | 0.269 | 11.353 | 509 |
| TCG | S | 0.125 | 11.286 | 506 |
| ACA | T | 0.264 | 11.13 | 499 |
| GGT | G | 0.218 | 10.394 | 466 |
| AGC | S | 0.115 | 10.372 | 465 |
| TGC | C | 0.449 | 9.903 | 444 |
| GCA | A | 0.309 | 9.837 | 441 |
| CTG | L | 0.094 | 9.747 | 437 |
| CAG | Q | 0.311 | 9.323 | 418 |
| GCT | A | 0.283 | 9.011 | 404 |
| GAC | D | 0.333 | 8.967 | 402 |
| GTG | V | 0.182 | 8.342 | 374 |
| GTC | V | 0.177 | 8.119 | 364 |
| CAC | H | 0.319 | 7.963 | 357 |
| CGG | R | 0.116 | 7.918 | 355 |
| CCG | P | 0.164 | 7.807 | 350 |
| GCC | A | 0.235 | 7.494 | 336 |
| GGC | G | 0.156 | 7.428 | 333 |
| ACG | T | 0.162 | 6.848 | 307 |
| CGT | R | 0.096 | 6.558 | 294 |
| CGC | R | 0.082 | 5.576 | 250 |
| GCG | A | 0.174 | 5.532 | 248 |

**Table S9. Statistics of the repeat elements in the Huizhan genome.**

| Type | Number of elements | Length (bp) | Percentage of genome (%) |
|---|---|---|---|
| Class I (Retrotransposons) | 103,839 | 114,027,649 | 28.85 |
| SINEs | 2,933 | 392,344 | 0.10 |
| LINEs | 11,690 | 5,472,942 | 1.38 |
| L1/CIN4 | 10,506 | 4,990,207 | 1.26 |
| LTR elements | 89,216 | 108,162,363 | 27.37 |
| Ty1/*Copia* | 19,084 | 14,382,692 | 3.64 |
| *Gypsy*/DIRS1 | 66,165 | 92,174,948 | 23.32 |
| Class II (DNA transposons) | 215,447 | 66,374,689 | 16.80 |
| hobo-Activator | 26,609 | 6,434,484 | 1.63 |
| Tc1-IS630-Pogo | 35,467 | 6,833,557 | 1.73 |
| Tourist/Harbinger | 45,417 | 10,446,831 | 2.64 |
| Other | 52 | 7,307 | 0.00 |
| Rolling-circles | 62,752 | 17,095,544 | 4.33 |
| Unclassified | 4,328 | 1,507,533 | 0.38 |
| Satellites | 449 | 5,873,648 | 1.49 |
| Total content | | 204,879,063 | 51.84 |

**Table S10. Functional annotation of the genes in Huizhan genome**

| | Tools/Database | Huizhan number | Huizhan percent (%) |
|---|---|---|---|
| Annotated | Interproscan | 35,158 | 0.89 |
| | eggNOG-mapper | 29,395 | 0.74 |
| | SwissProt | 19,937 | 0.50 |
| | NR | 36,046 | 0.91 |
| | KEGG | 8,403 | 0.21 |
| Total | | 38,157 | 96.46 |
| Unannotated | | 1,401 | 3.54 |

**Table S11. Statistics of the syntenic blocks between Huizhan and six other common XI rice genomes.**

| | Syntenic blocks | Self-sequence length (bp) | Huizhan-sequence length (bp) |
|---|---|---|---|
| IR64 | 13,938 | 360,947,993 | 361,020,060 |
| MH63 | 12,413 | 373,129,911 | 373,096,922 |
| ZS97 | 17,303 | 360,470,416 | 360,455,100 |
| R498 | 9,998 | 373,938,648 | 373,950,140 |
| Y58S | 14,267 | 364,594,734 | 364,586,687 |
| 93-11 | 12,780 | 363,417,981 | 363,447,571 |

**Table S12. Distribution of SNPs and InDels between Huizhan and six other common XI rice genomes.**

| | Type | IR64 | MH63 | ZS97 | R498 | Y58S | 93-11 |
|---|---|---|---|---|---|---|---|
| | Downstream | 146,239 | 128,415 | 171,953 | 103,255 | 157,099 | 131,508 |
| | Exonic | 88,017 | 84,479 | 102,945 | 65,068 | 95,650 | 83,589 |
| | Exonic; splicing | 0 | 1 | 1 | 2 | 2 | 2 |
| | Intergenic | 859,884 | 704,687 | 1,063,778 | 534,876 | 793,427 | 721,650 |
| | Intronic | 159,729 | 145,993 | 184,421 | 119,688 | 171,398 | 149,087 |
| SNPs | Splicing | 307 | 241 | 344 | 204 | 298 | 254 |
| | Upstream | 185,115 | 164,315 | 213,022 | 133,964 | 203,769 | 169,584 |
| | Upstream; downstream | 51,760 | 46,736 | 58,227 | 40,218 | 59,084 | 46,096 |
| | UTR3 | 23,558 | 20,807 | 26,130 | 17,111 | 26,294 | 21,684 |
| | UTR5 | 13,918 | 12,205 | 15,913 | 10,333 | 15,495 | 12,961 |
| | UTR5; UTR3 | 82 | 107 | 98 | 110 | 157 | 65 |
| | Downstream | 36,391 | 29,450 | 37,607 | 26,095 | 36,497 | 34,417 |
| | Exonic | 10,300 | 7,953 | 9,409 | 7,460 | 9,713 | 9,165 |
| | Exonic; splicing | 1 | 1 | 1 | 1 | 0 | 1 |
| | Intergenic | 178,982 | 114,588 | 155,580 | 111,837 | 142,782 | 144,162 |
| | Intronic | 46,332 | 39,187 | 47,047 | 35,358 | 46,187 | 44,862 |
| InDels | Splicing | 116 | 95 | 97 | 81 | 111 | 92 |
| | Upstream | 47,297 | 38,962 | 47,884 | 34,166 | 48,447 | 44,567 |
| | Upstream; downstream | 13,730 | 12,124 | 14,387 | 10,663 | 14,763 | 13,151 |
| | UTR3 | 7,333 | 6,471 | 7,967 | 5,490 | 8,091 | 7,259 |
| | UTR5 | 6,092 | 5,372 | 6,495 | 4,703 | 6,649 | 5,897 |
| | UTR5; UTR3 | 31 | 37 | 36 | 44 | 73 | 35 |

**Table S13. Functional annotations of genes involved in the inverted region of Chromosome 12**

| Gene ID | Function from Swissport |
|---|---|
| OsHZ37367 | RING-H2 finger protein |
| OsHZ37370 | Probable serine/threonine-protein kinase |
| OsHZ37372 | Pentatricopeptide repeat-containing protein |
| OsHZ37373 | Non-specific lipid-transfer protein |
| OsHZ37374 | Non-specific lipid-transfer protein |
| OsHZ37375 | Non-specific lipid-transfer protein |
| OsHZ37376 | Non-specific lipid-transfer protein |
| OsHZ37377 | Non-specific lipid-transfer protein |
| OsHZ37378 | Probable non-specific lipid-transfer protein |
| OsHZ37379 | RING-H2 finger protein |
| OsHZ37380 | Probable chalcone-flavonone isomerase |
| OsHZ37382 | Protein TOC75 |
| OsHZ37383 | Vacuolar sorting receptor |
| OsHZ37384 | Probable WRKY transcription factor |
| OsHZ37385 | WRKY transcription factor |
| OsHZ37387 | Probable WRKY transcription factor |
| OsHZ37388 | Probable WRKY transcription factor |
| OsHZ37391 | Methylesterase |
| OsHZ37392 | Root phototropism protein |
| OsHZ37393 | BTB/POZ domain-containing protein |
| OsHZ37402 | Ankyrin repeat domain-containing protein |
| OsHZ37404 | Cytochrome P450 |
| OsHZ37405 | Ankyrin repeat domain-containing protein |
| OsHZ37411 | Protein LURP-one-related |
| OsHZ37413 | Protein LURP-one-related |
| OsHZ37415 | Protein LURP-one-related |
| OsHZ37417 | Serine/threonine-protein kinase |
| OsHZ37418 | Ethanolamine-phosphate cytidylyltransferase |
| OsHZ37419 | Vacuolar protein sorting-associated protein |

**Table S14. The accession numbers of the 39 *Pi* genes and 15 *Xa* genes in this study**

| *Pi* genes | Accession numbers | *Xa* genes | Accession numbers |
|---|---|---|---|
| *Pi1-5* | AEB00617.1 | *Xa1* | BAA25068.1 |
| *Pi1-6* | AEB00618.1 | *Xa2/xa31* | QLH55888.1 |
| *Pi2* | ABC94599.1 | *Xa3/Xa26* | ABD84047.1 |
| *Pi21* | BAG72122.1 | *Xa4* | AQQ72921.1 |
| *Pi36* | ABI64281.1 | *Xa5* | AHC94892.1 |
| *Pi37* | ABI94578.1 | *Xa7* | QQY97222.1 |
| *Pi5-1* | ACJ54697.1 | *Xa10* | AGE45112.1 |
| *Pi5-2* | ACJ54698.1 | *Xa13* | ABD78942.1 |
| *Pi50_NBS4_3* | AKS24976.1 | *XA14* | QLH55889.1 |
| *Pi50_NBS4_1* | AKS24975.1 | *Xa21* | AAC49123.1 |
| *Pi7-1* | AET36551.1 | *Xa23* | AIX09984.1 |
| *Pi7-2* | AET36552.1 | *Xa25* | AGS56390.1 |
| *Pi9* | ABB88855.1 | *Xa27* | AAY54165.1 |
| *Pia* | BAK39922.1 | *Xa41* | NP_001410317.1 |
| *Pib* | BAA76281.2 | *xa45* | QLE11201.1 |
| *Pigm-R6* | APF29096.2 | | |
| *Pii-1* | QDZ58248.1 | | |
| *Pii-2* | QDZ58249.1 | | |
| *Pik-h* | AAY33493.1 | | |
| *Pikp-1* | ADV58352.1 | | |
| *Pikp-2* | ADV58351.1 | | |
| *Pik-1* | ADZ48537.1 | | |
| *Pik-2* | ADZ48538.1 | | |
| *Pikm1* | BAG72135.1 | | |
| *Pikm2* | BAG72136.1 | | |
| *Piks-1* | AET36547.1 | | |
| *Piks-2* | AET36548.1 | | |
| *Pish* | LOC_Os01g57340.1 | | |
| *Pit* | BAH20861.1 | | |
| *Pita* | AAK00132.1 | | |
| *Piz-t* | ABC73398.1 | | |
| *Pizh* | QBA09461.1 | | |
| *Ptr* | AWV95396.1 | | |
| *OsRGA5* | BAK39930.1 | | |
| *Pid3* | ACN79513.1 | | |
| *Pi63* | BAO79825.1 | | |
| *Pi56* | LOC_Os09g16000.1 | | |
| *Pi54* | CCD33072.1 | | |
| *Pb1* | AB570371 | | |

**Table S15. Statistics of the mapping rate of RNA-Seq data in Huizhan and ZH11 within heat stress**

| Type | Obtained Reads | OBtained Base (bp) | Q20 (%) | Q30 (%) | GC (%) | Mapping rate (%) | |
|---|---|---|---|---|---|---|---|
| | | | | | | Huizhan | Zhonghau 11 |
| Huizhan_flower_rep1 | 42,916,532 | 6,423,049,024 | 97.02 | 92.32 | 55.54 | 93.16 | - |
| Huizhan_flower_rep2 | 45,159,350 | 6,759,465,830 | 97.13 | 92.55 | 54.96 | 93.44 | - |
| Huizhan_flower_rep3 | 41,255,790 | 6,172,789,050 | 96.84 | 92.17 | 55.14 | 92.64 | - |
| ZH11_flower_rep1 | 40,214,840 | 6,016,880,298 | 96.68 | 91.73 | 54.73 | - | 95.18 |
| ZH11_flower_rep2 | 44,542,528 | 6,664,988,476 | 96.70 | 91.84 | 54.29 | - | 95.08 |
| ZH11_flower_rep3 | 45,085,884 | 6,746,735,602 | 96.91 | 92.27 | 53.43 | - | 95.38 |
| Huizhan_leaf_rep1 | 43,791,732 | 6,555,102,310 | 97.60 | 93.57 | 53.94 | 93.63 | - |
| Huizhan_leaf_rep2 | 39,355,880 | 5,888,745,890 | 96.70 | 91.70 | 53.90 | 92.39 | - |
| Huizhan_leaf_rep3 | 41,471,110 | 6,207,996,612 | 97.62 | 93.64 | 54.08 | 93.64 | - |
| ZH11_leaf_rep1 | 44,097,682 | 6,596,236,412 | 97.66 | 93.76 | 53.82 | - | 96.35 |
| ZH11_leaf_rep2 | 43,536,470 | 6,518,011,538 | 97.41 | 93.06 | 54.00 | - | 95.93 |
| ZH11_leaf_rep3 | 44,437,716 | 6,650,893,254 | 97.36 | 92.92 | 53.70 | - | 96.22 |

**Table S16. The GO enrichment of the up-regulated genes in flower group**

| GO acc | Description | Term | Ratio in foreground list | Ratio in background list | *p*-value | FDR | Number in foreground list |
|---|---|---|---|---|---|---|---|
| GO:0009737 | response to abscisic acid | BP | 67/1749 | 354/22598 | 6.29E-12 | 6.15E-09 | 67 |
| GO:0004568 | chitinase activity | MF | 19/1908 | 52/23764 | 8.18E-09 | 4.87E-06 | 19 |
| GO:0043565 | sequence-specific DNA binding | MF | 98/1908 | 706/23764 | 6.37E-08 | 1.90E-05 | 98 |
| GO:0010200 | response to chitin | BP | 34/1749 | 163/22598 | 8.62E-08 | 4.22E-05 | 34 |
| GO:0004857 | enzyme inhibitor activity | MF | 17/1908 | 52/23764 | 3.22E-07 | 6.41E-05 | 17 |
| GO:0009873 | ethylene-activated signaling pathway | BP | 36/1749 | 188/22598 | 3.44E-07 | 9.67E-05 | 36 |
| GO:0009414 | response to water deprivation | BP | 48/1749 | 289/22598 | 3.96E-07 | 9.67E-05 | 48 |
| GO:0071456 | cellular response to hypoxia | BP | 29/1749 | 149/22598 | 3.29E-06 | 0.000529 | 29 |
| GO:0050832 | defense response to fungus | BP | 54/1749 | 367/22598 | 3.74E-06 | 0.000529 | 54 |
| GO:0009611 | response to wounding | BP | 31/1749 | 166/22598 | 3.79E-06 | 0.000529 | 31 |
| GO:0042659 | regulation of cell fate specification | BP | 6/1749 | 8/22598 | 5.21E-06 | 0.000637 | 6 |
| GO:0045543 | gibberellin 2-beta-dioxygenase activity | MF | 6/1908 | 8/23764 | 6.46E-06 | 0.000662 | 6 |
| GO:0015333 | peptide:proton symporter activity | MF | 20/1908 | 83/23764 | 6.66E-06 | 0.000662 | 20 |
| GO:1904680 | peptide transmembrane transporter activity | MF | 20/1908 | 83/23764 | 6.66E-06 | 0.000662 | 20 |
| GO:0046658 | anchored component of plasma membrane | CC | 35/1797 | 191/21660 | 6.84E-06 | 0.000927 | 35 |
| GO:0005576 | extracellular region | CC | 138/1797 | 1158/21660 | 8.09E-06 | 0.000927 | 138 |
| GO:0005506 | iron ion binding | MF | 64/1908 | 470/23764 | 2.20E-05 | 0.0018 | 64 |
| GO:0034605 | cellular response to heat | BP | 14/1749 | 50/22598 | 1.73E-05 | 0.00188 | 14 |
| GO:0009625 | response to insect | BP | 9/1749 | 22/22598 | 1.92E-05 | 0.00188 | 9 |
| GO:0005975 | carbohydrate metabolic process | BP | 49/1749 | 345/22598 | 2.67E-05 | 0.00238 | 49 |
| GO:2000022 | regulation of jasmonic acid mediated signaling pathway | BP | 15/1749 | 59/22598 | 3.09E-05 | 0.0025 | 15 |

| | | | | | | | |
|---|---|---|---|---|---|---|---|
| GO:0045487 | gibberellin catabolic process | BP | 7/1749 | 14/22598 | 3.47E-05 | 0.0025 | 7 |
| GO:0048527 | lateral root development | BP | 15/1749 | 60/22598 | 3.82E-05 | 0.0025 | 15 |
| GO:0009738 | abscisic acid-activated signaling pathway | BP | 35/1749 | 222/22598 | 4.37E-05 | 0.00267 | 35 |
| GO:0020037 | heme binding | MF | 75/1908 | 591/23764 | 5.22E-05 | 0.00345 | 75 |
| GO:0015804 | neutral amino acid transport | BP | 6/1749 | 11/22598 | 7.00E-05 | 0.00381 | 6 |
| GO:0010333 | terpene synthase activity | MF | 19/1908 | 90/23764 | 8.03E-05 | 0.00467 | 19 |
| GO:0015171 | amino acid transmembrane transporter activity | MF | 18/1908 | 84/23764 | 1.00E-04 | 0.00497 | 18 |
| GO:0009409 | response to cold | BP | 41/1749 | 288/22598 | 0.000111 | 0.00543 | 41 |
| GO:0061408 | positive regulation of transcription from RNA polymerase II promoter in response to heat stress | BP | 9/1749 | 27/22598 | 0.000127 | 0.00564 | 9 |
| GO:0009636 | response to toxic substance | BP | 10/1749 | 33/22598 | 0.000133 | 0.00565 | 10 |
| GO:1900140 | regulation of seedling development | BP | 4/1749 | 5/22598 | 0.000168 | 0.00681 | 4 |
| GO:0043531 | ADP binding | MF | 72/1908 | 584/23764 | 0.000178 | 0.00809 | 72 |
| GO:0034485 | phosphatidylinositol-3,4,5-trisphosphate 5-phosphatase activity | MF | 7/1908 | 17/23764 | 0.000201 | 0.00809 | 7 |
| GO:0004497 | monooxygenase activity | MF | 36/1908 | 240/23764 | 0.000204 | 0.00809 | 36 |
| GO:0110126 | phloem loading | BP | 5/1749 | 9/22598 | 0.000267 | 0.00967 | 5 |
| GO:0008194 | UDP-glycosyltransferase activity | MF | 30/1908 | 190/23764 | 0.000273 | 0.0102 | 30 |
| GO:0051213 | dioxygenase activity | MF | 21/1908 | 115/23764 | 0.000303 | 0.0106 | 21 |
| GO:0004185 | serine-type carboxypeptidase activity | MF | 14/1908 | 62/23764 | 0.000325 | 0.0108 | 14 |
| GO:0042973 | glucan endo-1,3-beta-D-glucosidase activity | MF | 15/1908 | 71/23764 | 0.000435 | 0.0136 | 15 |
| GO:0001228 | DNA-binding transcription activator activity, RNA polymerase II-specific | MF | 10/1908 | 37/23764 | 0.000505 | 0.015 | 10 |
| GO:0018685 | alkane 1-monooxygenase activity | MF | 4/1908 | 6/23764 | 0.000545 | 0.0155 | 4 |
| GO:0010500 | transmitting tissue development | BP | 4/1749 | 6/22598 | 0.000472 | 0.0165 | 4 |

| | | | | | | | |
|---|---|---|---|---|---|---|---|
| GO:0048354 | mucilage biosynthetic process involved in seed coat development | BP | 7/1749 | 20/22598 | 0.000518 | 0.0175 | 7 |
| GO:0010336 | gibberellic acid homeostasis | BP | 5/1749 | 11/22598 | 0.000858 | 0.0249 | 5 |
| GO:0009685 | gibberellin metabolic process | BP | 6/1749 | 16/22598 | 0.000866 | 0.0249 | 6 |
| GO:0009646 | response to absence of light | BP | 9/1749 | 34/22598 | 0.000866 | 0.0249 | 9 |
| GO:0031640 | killing of cells of other organism | BP | 7/1749 | 22/22598 | 0.000993 | 0.027 | 7 |
| GO:0000272 | polysaccharide catabolic process | BP | 11/1749 | 51/22598 | 0.00153 | 0.0405 | 11 |
| GO:0015706 | nitrate transport | BP | 10/1749 | 44/22598 | 0.00163 | 0.0409 | 10 |
| GO:0016102 | diterpenoid biosynthetic process | BP | 10/1749 | 44/22598 | 0.00163 | 0.0409 | 10 |
| GO:0009688 | abscisic acid biosynthetic process | BP | 6/1749 | 18/22598 | 0.00175 | 0.0415 | 6 |
| GO:0009651 | response to salt stress | BP | 45/1749 | 371/22598 | 0.00182 | 0.0415 | 45 |
| GO:0007205 | protein kinase C-activating G protein-coupled receptor signaling pathway | BP | 4/1749 | 8/22598 | 0.00194 | 0.0415 | 4 |
| GO:0080168 | abscisic acid transport | BP | 4/1749 | 8/22598 | 0.00194 | 0.0415 | 4 |
| GO:0031098 | stress-activated protein kinase signaling cascade | BP | 10/1749 | 45/22598 | 0.00195 | 0.0415 | 10 |
| GO:0009627 | systemic acquired resistance | BP | 9/1749 | 38/22598 | 0.00203 | 0.0423 | 9 |
| GO:0050502 | cis-zeatin O-beta-D-glucosyltransferase activity | MF | 5/1908 | 12/23764 | 0.00163 | 0.0428 | 5 |
| GO:0051119 | sugar transmembrane transporter activity | MF | 7/1908 | 23/23764 | 0.00165 | 0.0428 | 7 |
| GO:0071555 | cell wall organization | BP | 45/1749 | 375/22598 | 0.00224 | 0.0449 | 45 |
| GO:0046856 | phosphatidylinositol dephosphorylation | BP | 7/1749 | 25/22598 | 0.00228 | 0.0449 | 7 |
| GO:0023014 | signal transduction by protein phosphorylation | BP | 10/1749 | 46/22598 | 0.00233 | 0.0449 | 10 |
| GO:0009813 | flavonoid biosynthetic process | BP | 15/1749 | 86/22598 | 0.00234 | 0.0449 | 15 |
| GO:0000978 | RNA polymerase II proximal promoter sequence-specific DNA binding | MF | 17/1908 | 98/23764 | 0.00196 | 0.0467 | 17 |
| GO:0006749 | glutathione metabolic process | BP | 14/1749 | 79/22598 | 0.00277 | 0.0492 | 14 |
| GO:1902584 | positive regulation of response to water deprivation | BP | 7/1749 | 26/22598 | 0.00291 | 0.0494 | 7 |

| | | | | | | | |
|---|---|---|---|---|---|---|---|
| GO:0006032 | chitin catabolic process | BP | 9/1749 | 40/22598 | 0.00296 | 0.0494 | 9 |
| GO:0042128 | nitrate assimilation | BP | 9/1749 | 40/22598 | 0.00296 | 0.0494 | 9 |
| GO:0009626 | plant-type hypersensitive response | BP | 30/1749 | 228/22598 | 0.003 | 0.0494 | 30 |
| GO:0005992 | trehalose biosynthetic process | BP | 6/1749 | 20/22598 | 0.0032 | 0.0494 | 6 |
| GO:0030001 | metal ion transport | BP | 10/1749 | 48/22598 | 0.00324 | 0.0494 | 10 |
| GO:0006809 | nitric oxide biosynthetic process | BP | 4/1749 | 9/22598 | 0.00328 | 0.0494 | 4 |
| GO:0048262 | determination of dorsal/ventral asymmetry | BP | 4/1749 | 9/22598 | 0.00328 | 0.0494 | 4 |
| GO:0048856 | anatomical structure development | BP | 4/1749 | 9/22598 | 0.00328 | 0.0494 | 4 |
| GO:0004439 | phosphatidylinositol-4,5-bisphosphate 5-phosphatase activity | MF | 7/1908 | 24/23764 | 0.00217 | 0.0498 | 7 |

**Table S17. The GO enrichment of the up-regulated genes in leaf group**

| GO acc | Description | Term | Ratio in foreground list | Ratio in background list | *p*-value | FDR | Number in foreground list |
|---|---|---|---|---|---|---|---|
| GO:0008194 | UDP-glycosyltransferase activity | MF | 34/1640 | 190/23764 | 2.69E-07 | 0.000159 | 34 |
| GO:0043531 | ADP binding | MF | 71/1640 | 584/23764 | 2.35E-06 | 0.000692 | 71 |
| GO:0015181 | arginine transmembrane transporter activity | MF | 6/1640 | 9/23764 | 7.50E-06 | 0.00114 | 6 |
| GO:0080044 | quercetin 7-O-glucosyltransferase activity | MF | 14/1640 | 52/23764 | 7.70E-06 | 0.00114 | 14 |
| GO:0016807 | cysteine-type carboxypeptidase activity | MF | 4/1640 | 5/23764 | 0.000107 | 0.00887 | 4 |
| GO:1990380 | Lys48-specific deubiquitinase activity | MF | 4/1640 | 5/23764 | 0.000107 | 0.00887 | 4 |
| GO:0080167 | response to karrikin | BP | 19/1483 | 94/22598 | 9.44E-06 | 0.00963 | 19 |
| GO:0005315 | inorganic phosphate transmembrane transporter activity | MF | 8/1640 | 27/23764 | 0.000343 | 0.0184 | 8 |
| GO:0022857 | transmembrane transporter activity | MF | 28/1640 | 204/23764 | 0.000388 | 0.0191 | 28 |
| GO:0004709 | MAP kinase kinase kinase activity | MF | 8/1640 | 28/23764 | 0.000451 | 0.0192 | 8 |
| GO:0030246 | carbohydrate binding | MF | 42/1640 | 355/23764 | 0.000456 | 0.0192 | 42 |
| GO:0052634 | C-19 gibberellin 2-beta-dioxygenase activity | MF | 5/1640 | 11/23764 | 0.000505 | 0.0199 | 5 |
| GO:0009626 | plant-type hypersensitive response | BP | 32/1483 | 228/22598 | 3.92E-05 | 0.02 | 32 |
| GO:0003979 | UDP-glucose 6-dehydrogenase activity | MF | 4/1640 | 7/23764 | 0.000668 | 0.0232 | 4 |
| GO:0004784 | superoxide dismutase activity | MF | 4/1640 | 7/23764 | 0.000668 | 0.0232 | 4 |
| GO:0034605 | cellular response to heat | BP | 12/1483 | 50/22598 | 7.16E-05 | 0.0244 | 12 |
| GO:0015297 | antiporter activity | MF | 19/1640 | 125/23764 | 0.000936 | 0.0307 | 19 |
| GO:0043069 | negative regulation of programmed cell death | BP | 5/1483 | 9/22598 | 0.000122 | 0.0311 | 5 |
| GO:0009611 | response to wounding | BP | 24/1483 | 166/22598 | 0.00022 | 0.0413 | 24 |
| GO:0061408 | positive regulation of transcription from RNA polymerase II promoter in response to heat stress | BP | 8/1483 | 27/22598 | 0.000243 | 0.0413 | 8 |

**Table S18. Statistics analysis of HSF genes within DEGs. Yellow represents upregulated genes, and blue represents downregulated genes.**

| Flower | HSF | Leaf | HSF |
|---|---|---|---|
| LOC_Os03g06630 | *HsfA2d* | LOC_Os01g54550 | *HsfA4b* |
| LOC_Os02g32590 | *HsfA3* | LOC_Os05g45410 | *HsfA4d* |
| LOC_Os01g39020 | *HsfA6b* | LOC_Os03g12370 | *HsfA9* |
| LOC_Os03g12370 | *HsfA9* | LOC_Os09g28354 | *HsfB1* |
| LOC_Os09g28354 | *HsfB1* | LOC_Os04g48030 | *HsfB2a* |
| LOC_Os01g43590 | *HsfC1a* | LOC_Os08g43334 | *HsfB2b* |
| LOC_Os01g53220 | *HsfC1b* | LOC_Os09g35790 | *HsfB2c* |
| LOC_Os02g13800 | *HsfC2a* | LOC_Os01g53220 | *HsfC1b* |
| LOC_Os06g35960 | *HsfC2b* | | |
| LOC_Os07g44690 | *HsfB4b* | | |

**Table S19. Statistics analysis of transcription factors NAC, WYB, and WRKY within DEGs. Yellow represents upregulated genes, and blue represents downregulated genes.**

| Flower | | | Leaf | | |
|---|---|---|---|---|---|
| NAC | WYB | WRKY | NAC | WYB | WRKY |
| LOC_Os01g15640 | LOC_Os01g18240 | LOC_Os01g09100 | LOC_Os01g47670 | LOC_Os01g45090 | LOC_Os01g14440 |
| LOC_Os01g64310 | LOC_Os01g50110 | LOC_Os01g14440 | LOC_Os01g66120 | LOC_Os01g50720 | LOC_Os01g40260 |
| LOC_Os01g66120 | LOC_Os01g63460 | LOC_Os01g43650 | LOC_Os02g18460 | LOC_Os01g64360 | LOC_Os01g43550 |
| LOC_Os02g36880 | LOC_Os01g64360 | LOC_Os01g47560 | LOC_Os02g38130 | LOC_Os02g09480 | LOC_Os01g46800 |
| LOC_Os02g41450 | LOC_Os01g74410 | LOC_Os01g53040 | LOC_Os03g42630 | LOC_Os02g41510 | LOC_Os01g47560 |
| LOC_Os02g56600 | LOC_Os02g40530 | LOC_Os01g54600 | LOC_Os05g37080 | LOC_Os02g42850 | LOC_Os01g53040 |
| LOC_Os03g01870 | LOC_Os02g42850 | LOC_Os01g60600 | LOC_Os06g01480 | LOC_Os05g37060 | LOC_Os02g08440 |
| LOC_Os03g21030 | LOC_Os02g46780 | LOC_Os01g60640 | LOC_Os06g36480 | LOC_Os05g37730 | LOC_Os02g16540 |
| LOC_Os03g56580 | LOC_Os02g54520 | LOC_Os02g08440 | LOC_Os10g21560 | LOC_Os06g40330 | LOC_Os03g33012 |
| LOC_Os03g60080 | LOC_Os03g04900 | LOC_Os03g20550 | LOC_Os10g27360 | LOC_Os06g43090 | LOC_Os03g55080 |
| LOC_Os04g35660 | LOC_Os03g20090 | LOC_Os03g55080 | LOC_Os10g42130 | LOC_Os08g33150 | LOC_Os05g50700 |
| LOC_Os04g38720 | LOC_Os03g26130 | LOC_Os04g21950 | LOC_Os11g05614 | LOC_Os09g23620 | LOC_Os06g44010 |
| LOC_Os04g43560 | LOC_Os04g43680 | LOC_Os05g25770 | LOC_Os12g04230 | LOC_Os01g09590 | LOC_Os07g40570 |
| LOC_Os05g10620 | LOC_Os05g37060 | LOC_Os05g27730 | LOC_Os12g41680 | LOC_Os01g19330 | LOC_Os11g45850 |
| LOC_Os05g34830 | LOC_Os06g14670 | LOC_Os05g49100 | LOC_Os03g04070 | LOC_Os02g40530 | LOC_Os11g45920 |
| LOC_Os05g37080 | LOC_Os06g43090 | LOC_Os05g50610 | LOC_Os03g21060 | LOC_Os03g51110 | LOC_Os01g09100 |
| LOC_Os06g04090 | LOC_Os06g46560 | LOC_Os05g50700 | LOC_Os04g38720 | LOC_Os05g48010 | LOC_Os01g60640 |
| LOC_Os06g36480 | LOC_Os07g48870 | LOC_Os06g05380 | LOC_Os07g37920 | LOC_Os07g43580 | LOC_Os05g03900 |
| LOC_Os06g46270 | LOC_Os09g01960 | LOC_Os09g25060 | LOC_Os08g10080 | LOC_Os08g43550 | LOC_Os05g25770 |
| LOC_Os07g48450 | LOC_Os11g03440 | LOC_Os11g29870 | LOC_Os08g33670 | LOC_Os09g36250 | LOC_Os05g46020 |

| | | | | | |
|---|---|---|---|---|---|
| LOC_Os07g48550 | LOC_Os02g42870 | LOC_Os11g45850 | LOC_Os09g33490 | LOC_Os11g35390 | LOC_Os05g49620 |
| LOC_Os08g01330 | LOC_Os09g36250 | LOC_Os11g45920 | LOC_Os11g03300 | LOC_Os11g45740 | LOC_Os07g48260 |
| LOC_Os08g02160 | | LOC_Os12g02420 | LOC_Os11g04960 | LOC_Os11g47460 | LOC_Os08g13840 |
| LOC_Os08g02300 | | LOC_Os08g13840 | LOC_Os12g03040 | LOC_Os12g37970 | LOC_Os08g29660 |
| LOC_Os08g42400 | | | | | LOC_Os09g16510 |
| LOC_Os09g33490 | | | | | LOC_Os11g02520 |
| LOC_Os10g38834 | | | | | LOC_Os11g02540 |
| LOC_Os10g42130 | | | | | |
| LOC_Os11g05614 | | | | | |
| LOC_Os11g08210 | | | | | |
| LOC_Os12g03050 | | | | | |
| LOC_Os12g22630 | | | | | |
| LOC_Os04g52810 | | | | | |
| LOC_Os08g10080 | | | | | |
| LOC_Os11g04960 | | | | | |

**Table S20. Correlation analyses of nine HSFs to other genes in flower group. Significant differences were tested by two-tailed Pearson correlation test and shown by *p*-value.**

| | LOC_Os06g35960 | LOC_Os02g32590 | LOC_Os01g39020 | LOC_Os01g53220 | LOC_Os02g13800 | LOC_Os01g43590 | LOC_Os03g12370 | LOC_Os03g06630 | LOC_Os09g28354 |
|---|---|---|---|---|---|---|---|---|---|
| NAC | | | | | | | | | |
| LOC_Os06g46270 | 0.9615835 | 0.6887934 | 0.1754224 | 0.4577453 | 0.848458 | 0.002193315 | 0.32288 | 0.3532185 | 0.05356207 |
| LOC_Os05g10620 | 0.9757469 | 0.6261238 | 0.238092 | 0.3950757 | 0.9111276 | 0.06486288 | 0.3855495 | 0.2905489 | 0.1162316 |
| LOC_Os10g42130 | 0.2840692 | 0.6336923 | 0.5020919 | 0.8647404 | 0.1709437 | 0.675321 | 0.3546343 | 0.9692672 | 0.6239522 |
| LOC_Os11g05614 | 0.3648652 | 0.01524209 | 0.8489738 | 0.2158061 | 0.4779907 | 0.6757447 | 0.9964313 | 0.3203329 | 0.7271134 |
| LOC_Os04g35660 | 0.8485898 | 0.4989667 | 0.3652492 | 0.2679185 | 0.9617153 | 0.19202 | 0.5127067 | 0.1633917 | 0.2433888 |
| LOC_Os02g56600 | 0.7490509 | 0.3994277 | 0.4647881 | 0.1683796 | 0.8621764 | 0.291559 | 0.6122456 | 0.0638528 | 0.3429277 |
| LOC_Os12g03050 | 0.4207788 | 0.7704018 | 0.3653823 | 0.99855 | 0.3076532 | 0.5386115 | 0.2179248 | 0.8940232 | 0.4872427 |
| LOC_Os10g38834 | 0.6495382 | 0.9991612 | 0.1366229 | 0.7697906 | 0.5364127 | 0.309852 | 0.01083465 | 0.6652638 | 0.2584833 |
| LOC_Os06g04090 | 0.4939335 | 0.8435566 | 0.2922276 | 0.9253953 | 0.380808 | 0.4654567 | 0.14477 | 0.8208685 | 0.414088 |

| | | | | | | | | | |
|---|---|---|---|---|---|---|---|---|---|
| LOC_Os04g38720 | 0.3138359 | 0.03578723 | 0.9000031 | 0.2668354 | 0.4269614 | 0.726774 | 0.9525394 | 0.3713622 | 0.7781427 |
| LOC_Os07g48450 | 0.9366071 | 0.7137699 | 0.150446 | 0.4827217 | 0.8234816 | 0.02278311 | 0.2979035 | 0.3781949 | 0.02858564 |
| LOC_Os08g02160 | 0.3184746 | 0.6680977 | 0.4676865 | 0.8991458 | 0.2053491 | 0.6409156 | 0.320229 | 0.9963274 | 0.5895469 |
| LOC_Os03g56580 | 0.8738685 | 0.5242454 | 0.3399705 | 0.2931972 | 0.986994 | 0.1667414 | 0.487428 | 0.1886704 | 0.2181101 |
| LOC_Os03g01870 | 0.639362 | 0.9889851 | 0.146799 | 0.7799667 | 0.5262365 | 0.3200281 | 0.000658526 | 0.6754399 | 0.2686594 |
| LOC_Os11g08210 | 0.8845429 | 0.765834 | 0.09838185 | 0.5347859 | 0.7714174 | 0.07484728 | 0.2458394 | 0.430259 | 0.02347853 |
| LOC_Os12g22630 | 0.6288524 | 0.2792293 | 0.5849866 | 0.04818112 | 0.7419779 | 0.4117574 | 0.7324441 | 0.05634567 | 0.4631262 |
| LOC_Os03g60080 | 0.7155471 | 0.9348298 | 0.07061393 | 0.7037816 | 0.6024216 | 0.2438431 | 0.0768436 | 0.5992548 | 0.1924743 |
| LOC_Os05g34830 | 0.9702502 | 0.6801267 | 0.1840892 | 0.4490785 | 0.8571247 | 0.01086007 | 0.3315467 | 0.3445517 | 0.06222882 |
| LOC_Os05g37080 | 0.3482602 | 0.6978833 | 0.4379008 | 0.9289315 | 0.2351347 | 0.6111299 | 0.2904433 | 0.9665417 | 0.5597612 |
| LOC_Os02g41450 | 0.3878384 | 0.03821535 | 0.8260005 | 0.1928328 | 0.5009639 | 0.6527714 | 0.9734581 | 0.2973596 | 0.7041401 |
| LOC_Os01g15640 | 0.6257591 | 0.276136 | 0.5880798 | 0.04508787 | 0.7388846 | 0.4148507 | 0.7355374 | 0.05943891 | 0.4662195 |
| LOC_Os08g02300 | 0.4276833 | 0.7773064 | 0.3584777 | 0.9916455 | 0.3145578 | 0.5317069 | 0.2110202 | 0.8871186 | 0.4803381 |

| | | | | | | | | | |
|---|---|---|---|---|---|---|---|---|---|
| LOC_Os04g43560 | 0.9454169 | 0.70496 | 0.1592558 | 0.4739119 | 0.8322914 | 0.01397331 | 0.3067133 | 0.3693851 | 0.03739544 |
| LOC_Os07g48550 | 0.6792322 | 0.9711447 | 0.1069288 | 0.7400965 | 0.5661067 | 0.2801579 | 0.04052873 | 0.6355697 | 0.2287892 |
| LOC_Os03g21030 | 0.007844511 | 0.3574676 | 0.7783166 | 0.5885158 | 0.105281 | 0.9515457 | 0.630859 | 0.6930425 | 0.9001769 |
| LOC_Os09g33490 | 0.7654498 | 0.8849272 | 0.02071127 | 0.653879 | 0.6523243 | 0.1939404 | 0.1267463 | 0.5493522 | 0.1425716 |
| LOC_Os02g36880 | 0.7746255 | 0.4250025 | 0.4392134 | 0.1939543 | 0.8877511 | 0.2659842 | 0.5866709 | 0.08942754 | 0.317353 |
| LOC_Os08g42400 | 0.6063057 | 0.2566825 | 0.6075333 | 0.0256344 | 0.7194312 | 0.4343042 | 0.7549908 | 0.07889239 | 0.4856729 |
| LOC_Os08g01330 | 0.89198 | 0.7583969 | 0.105819 | 0.5273488 | 0.7788545 | 0.06741017 | 0.2532765 | 0.4228219 | 0.01604142 |
| LOC_Os01g66120 | 0.1575758 | 0.1920473 | 0.9437369 | 0.4230955 | 0.2707013 | 0.883034 | 0.7962793 | 0.5276222 | 0.9344028 |
| LOC_Os06g36480 | 0.353715 | 0.7033381 | 0.4324461 | 0.9343862 | 0.2405895 | 0.6056752 | 0.2849886 | 0.961087 | 0.5543064 |
| LOC_Os01g64310 | 0.6310433 | 0.9806663 | 0.1551178 | 0.7882855 | 0.5179178 | 0.3283469 | 0.007660258 | 0.6837587 | 0.2769782 |
| WYB | | | | | | | | | |
| LOC_Os04g43680 | 0.3942854 | 0.7439085 | 0.3918757 | 0.9749566 | 0.2811599 | 0.5651048 | 0.2444182 | 0.9205166 | 0.5137361 |
| LOC_Os05g37060 | 0.6079227 | 0.9575458 | 0.1782384 | 0.8114061 | 0.4947971 | 0.3514675 | 0.03078089 | 0.7068793 | 0.3000988 |

| | | | | | | | | | |
|---|---|---|---|---|---|---|---|---|---|
| LOC_Os02g42850 | 0.5264384 | 0.8760614 | 0.2597227 | 0.8928904 | 0.4133129 | 0.4329518 | 0.1122652 | 0.7883636 | 0.3815831 |
| LOC_Os03g20090 | 0.6806778 | 0.9696991 | 0.1054833 | 0.738651 | 0.5675523 | 0.2787124 | 0.04197426 | 0.6341242 | 0.2273436 |
| LOC_Os06g14670 | 0.2202743 | 0.1293488 | 0.9935647 | 0.360397 | 0.3333998 | 0.8203355 | 0.8589779 | 0.4649237 | 0.8717043 |
| LOC_Os02g46780 | 0.9324637 | 0.7179132 | 0.1463027 | 0.486865 | 0.8193382 | 0.02692648 | 0.2937602 | 0.3823383 | 0.02444228 |
| LOC_Os03g04900 | 0.794177 | 0.8561999 | 0.008015916 | 0.6251518 | 0.6810515 | 0.1652132 | 0.1554734 | 0.520625 | 0.1138445 |
| LOC_Os01g50110 | 0.9794381 | 0.629815 | 0.2344008 | 0.3987669 | 0.9074363 | 0.06117166 | 0.3818583 | 0.2942401 | 0.1125404 |
| LOC_Os02g40530 | 0.6072977 | 0.9569208 | 0.1788633 | 0.812031 | 0.4941722 | 0.3520925 | 0.03140581 | 0.7075043 | 0.3007237 |
| LOC_Os01g18240 | 0.6396479 | 0.989271 | 0.1465132 | 0.7796809 | 0.5265223 | 0.3197423 | 0.000944338 | 0.6751541 | 0.2683736 |
| LOC_Os02g54520 | 0.7334203 | 0.3837972 | 0.4804187 | 0.152749 | 0.8465458 | 0.3071895 | 0.6278762 | 0.04822226 | 0.3585583 |
| LOC_Os11g03440 | 0.8041716 | 0.8462053 | 0.01801057 | 0.6151571 | 0.6910461 | 0.1552186 | 0.1654681 | 0.5106304 | 0.1038498 |
| LOC_Os01g63460 | 0.9877848 | 0.6381617 | 0.2260541 | 0.4071136 | 0.8990897 | 0.052825 | 0.3735117 | 0.3025868 | 0.1041938 |
| LOC_Os06g43090 | 0.7309764 | 0.9194005 | 0.05518463 | 0.6883523 | 0.6178509 | 0.2284138 | 0.09227289 | 0.5838255 | 0.177045 |
| LOC_Os01g64360 | 0.7990264 | 0.8513505 | 0.01286537 | 0.6203023 | 0.6859009 | 0.1603638 | 0.1603229 | 0.5157756 | 0.108995 |

| | | | | | | | | | |
|---|---|---|---|---|---|---|---|---|---|
| LOC_Os07g48870 | 0.8558161 | 0.506193 | 0.3580228 | 0.2751449 | 0.9689416 | 0.1847937 | 0.5054803 | 0.1706181 | 0.2361625 |
| LOC_Os06g46560 | 0.7631078 | 0.4134847 | 0.4507312 | 0.1824365 | 0.8762333 | 0.2775021 | 0.5981887 | 0.07790972 | 0.3288708 |
| LOC_Os09g01960 | 0.9926917 | 0.6576852 | 0.2065306 | 0.4266371 | 0.8795662 | 0.0333015 | 0.3539881 | 0.3221103 | 0.08467025 |
| LOC_Os03g26130 | 0.8410332 | 0.8093437 | 0.05487214 | 0.5782955 | 0.7279077 | 0.118357 | 0.2023297 | 0.4737688 | 0.06698824 |
| LOC_Os01g74410 | 0.625304 | 0.9749271 | 0.1608571 | 0.7940248 | 0.5121784 | 0.3340862 | 0.01339958 | 0.689498 | 0.2827175 |
| WRKY | | | | | | | | | |
| LOC_Os11g45920 | 0.5291598 | 0.8787829 | 0.2570013 | 0.890169 | 0.4160343 | 0.4302304 | 0.1095438 | 0.7856422 | 0.3788617 |
| LOC_Os01g60600 | 0.9711103 | 0.6792666 | 0.1849493 | 0.4482184 | 0.8579848 | 0.01172016 | 0.3324068 | 0.3436916 | 0.06308891 |
| LOC_Os04g21950 | 0.4785751 | 0.1289521 | 0.7352638 | 0.1020961 | 0.5917007 | 0.5620347 | 0.8827213 | 0.2066229 | 0.6134034 |
| LOC_Os05g50700 | 0.2333027 | 0.5829257 | 0.5528584 | 0.8139739 | 0.1201772 | 0.7260875 | 0.4054009 | 0.9185007 | 0.6747188 |
| LOC_Os11g29870 | 0.7501066 | 0.4004835 | 0.4637323 | 0.1694354 | 0.8632321 | 0.2905032 | 0.6111898 | 0.06490859 | 0.3418719 |
| LOC_Os05g49100 | 0.9664209 | 0.6167978 | 0.2474181 | 0.3857496 | 0.9204536 | 0.07418895 | 0.3948756 | 0.2812229 | 0.1255577 |
| LOC_Os03g55080 | 0.4000219 | 0.749645 | 0.3861391 | 0.9806932 | 0.2868964 | 0.5593683 | 0.2386816 | 0.91478 | 0.5079995 |

| | | | | | | | | | |
|---|---|---|---|---|---|---|---|---|---|
| LOC_Os01g14440 | 0.1057438 | 0.4553669 | 0.6804172 | 0.6864151 | 0.007381699 | 0.8536464 | 0.5329597 | 0.7909418 | 0.8022776 |
| LOC_Os09g25060 | 0.3634739 | 0.713097 | 0.4226872 | 0.9441451 | 0.2503484 | 0.5959163 | 0.2752297 | 0.9513281 | 0.5445476 |
| LOC_Os03g20550 | 0.9548437 | 0.6955332 | 0.1686826 | 0.464485 | 0.8417182 | 0.004546496 | 0.3161401 | 0.3599583 | 0.04682226 |
| LOC_Os01g54600 | 0.7569681 | 0.8934088 | 0.02919297 | 0.6623607 | 0.6438426 | 0.2024221 | 0.1182646 | 0.5578339 | 0.1510534 |
| LOC_Os05g27730 | 0.3662199 | 0.715843 | 0.4199411 | 0.9468912 | 0.2530944 | 0.5931702 | 0.2724836 | 0.9485821 | 0.5418015 |
| LOC_Os01g53040 | 0.8901943 | 0.5405712 | 0.3236447 | 0.309523 | 0.9966802 | 0.1504155 | 0.4711022 | 0.2049962 | 0.2017843 |
| LOC_Os05g50610 | 0.1536949 | 0.503318 | 0.6324661 | 0.7343661 | 0.04056939 | 0.8056953 | 0.4850086 | 0.8388929 | 0.7543265 |
| LOC_Os02g08440 | 0.3873475 | 0.7369706 | 0.3988136 | 0.9680187 | 0.274222 | 0.5720427 | 0.251356 | 0.9274545 | 0.5206739 |
| LOC_Os05g25770 | 0.5397434 | 0.8893665 | 0.2464177 | 0.8795854 | 0.4266179 | 0.4196468 | 0.09896013 | 0.7750586 | 0.368278 |
| LOC_Os06g05380 | 0.8177562 | 0.8326207 | 0.03159509 | 0.6015726 | 0.7046306 | 0.141634 | 0.1790526 | 0.4970458 | 0.09026529 |
| LOC_Os11g45850 | 0.8688195 | 0.7815574 | 0.08265848 | 0.5505092 | 0.755694 | 0.09057065 | 0.230116 | 0.4459824 | 0.0392019 |
| LOC_Os01g09100 | 0.7600671 | 0.8903098 | 0.02609392 | 0.6592616 | 0.6469416 | 0.1993231 | 0.1213636 | 0.5547348 | 0.1479543 |
| LOC_Os12g02420 | 0.3384027 | 0.6880258 | 0.4477583 | 0.919074 | 0.2252772 | 0.6209875 | 0.3003008 | 0.9763992 | 0.5696187 |

| | | | | | | | | | |
|---|---|---|---|---|---|---|---|---|---|
| LOC_Os01g43650 | 0.5131722 | 0.8627953 | 0.2729888 | 0.9061565 | 0.4000467 | 0.446218 | 0.1255313 | 0.8016297 | 0.3948492 |
| LOC_Os01g47560 | 0.4941559 | 0.1445329 | 0.719683 | 0.0865153 | 0.6072814 | 0.5464539 | 0.8671405 | 0.1910421 | 0.5978226 |
| LOC_Os01g60640 | 0.3975055 | 0.04788242 | 0.8163335 | 0.1831657 | 0.510631 | 0.6431043 | 0.963791 | 0.2876925 | 0.6944731 |

**Table S21. Correlation analyses of eight HSFs to other genes in leaf group. Significant differences were tested by two-tailed Pearson correlation test and shown by *p*-value.**

| | LOC_Os01g54550 | LOC_Os01g53220 | LOC_Os08g43334 | LOC_Os09g35790 | LOC_Os05g45410 | LOC_Os04g48030 | LOC_Os03g12370 | LOC_Os09g28354 |
|---|---|---|---|---|---|---|---|---|
| NAC | | | | | | | | |
| LOC_Os10g42130 | 0.4671986 | 0.8647404 | 0.2128253 | 0.9575154 | 0.624339 | 0.3113329 | 0.3546343 | 0.6239522 |
| LOC_Os11g05614 | 0.883867 | 0.2158061 | 0.4361091 | 0.3935503 | 0.02459535 | 0.9602673 | 0.9964313 | 0.7271134 |
| LOC_Os01g47670 | 0.5324836 | 0.7994555 | 0.1475403 | 0.9771997 | 0.5590541 | 0.3766179 | 0.4199193 | 0.6892372 |
| LOC_Os10g27360 | 0.9958055 | 0.3361335 | 0.3157816 | 0.5138777 | 0.09573212 | 0.8399398 | 0.8832412 | 0.8474408 |
| LOC_Os02g18460 | 0.2138127 | 0.8818737 | 0.4662112 | 0.7041295 | 0.8777249 | 0.05794702 | 0.1012484 | 0.3705663 |
| LOC_Os06g01480 | 0.3069389 | 0.361122 | 0.9869628 | 0.1833778 | 0.6015235 | 0.4628046 | 0.4195032 | 0.1501853 |
| LOC_Os12g41680 | 0.4376698 | 0.2303911 | 0.8823063 | 0.05264691 | 0.4707925 | 0.5935355 | 0.5502341 | 0.2809162 |
| LOC_Os05g37080 | 0.4030076 | 0.9289315 | 0.2770163 | 0.8933243 | 0.6885301 | 0.2471419 | 0.2904433 | 0.5597612 |
| LOC_Os10g21560 | 0.3940781 | 0.937861 | 0.2859458 | 0.8843948 | 0.6974595 | 0.2382124 | 0.2815138 | 0.5508317 |

| | | | | | | | | |
|---|---|---|---|---|---|---|---|---|
| LOC_Os03g42630 | 0.7007849 | 0.6311541 | 0.02076102 | 0.8088983 | 0.3907527 | 0.5449192 | 0.5882207 | 0.8575385 |
| LOC_Os02g38130 | 0.1046837 | 0.7727446 | 0.5753402 | 0.5950004 | 0.986854 | 0.05118203 | 0.00788062 | 0.2614373 |
| LOC_Os01g66120 | 0.9088436 | 0.4230955 | 0.2288197 | 0.6008397 | 0.182694 | 0.7529779 | 0.7962793 | 0.9344028 |
| LOC_Os06g36480 | 0.3975528 | 0.9343862 | 0.2824711 | 0.8878696 | 0.6939848 | 0.2416871 | 0.2849886 | 0.5543064 |
| LOC_Os12g04230 | 0.5674098 | 0.7645293 | 0.1126141 | 0.9422735 | 0.5241279 | 0.4115441 | 0.4548455 | 0.7241634 |
| WYB | | | | | | | | |
| LOC_Os05g37060 | 0.1433452 | 0.8114061 | 0.5366787 | 0.6336619 | 0.9481925 | 0.01252053 | 0.03078089 | 0.3000988 |
| LOC_Os02g42850 | 0.2248295 | 0.8928904 | 0.4551944 | 0.7151462 | 0.8667082 | 0.06896377 | 0.1122652 | 0.3815831 |
| LOC_Os02g09480 | 0.1951004 | 0.8631613 | 0.4849235 | 0.6854171 | 0.8964372 | 0.0392347 | 0.08253611 | 0.351854 |
| LOC_Os09g23620 | 0.8586421 | 0.4732969 | 0.1786182 | 0.6510412 | 0.2328955 | 0.7027764 | 0.7460778 | 0.9846042 |
| LOC_Os06g40330 | 0.4618986 | 0.2061624 | 0.8580776 | 0.02841819 | 0.4465638 | 0.6177642 | 0.5744628 | 0.3051449 |
| LOC_Os06g43090 | 0.02029138 | 0.6883523 | 0.6597325 | 0.5106081 | 0.9287537 | 0.1355743 | 0.09227289 | 0.177045 |
| LOC_Os01g64360 | 0.04775862 | 0.6203023 | 0.7277825 | 0.4425581 | 0.8607038 | 0.2036243 | 0.1603229 | 0.108995 |

| | | | | | | | | |
|---|---|---|---|---|---|---|---|---|
| LOC_Os01g45090 | 0.3177105 | 0.3503505 | 0.9977344 | 0.1726063 | 0.5907519 | 0.4735762 | 0.4302748 | 0.1609569 |
| LOC_Os08g33150 | 0.2799702 | 0.9480311 | 0.4000537 | 0.7702869 | 0.8115675 | 0.1241045 | 0.1674059 | 0.4367238 |
| LOC_Os01g50720 | 0.07456902 | 0.5934919 | 0.7545929 | 0.4157477 | 0.8338934 | 0.2304347 | 0.1871333 | 0.0821846 |
| LOC_Os02g41510 | 0.07665317 | 0.5914078 | 0.756677 | 0.4136636 | 0.8318092 | 0.2325189 | 0.1892174 | 0.08010045 |
| LOC_Os05g37730 | 0.4371178 | 0.8948212 | 0.242906 | 0.9274346 | 0.6544198 | 0.2812522 | 0.3245536 | 0.5938715 |
| WRKY | | | | | | | | |
| LOC_Os11g45920 | 0.2221081 | 0.890169 | 0.4579158 | 0.7124248 | 0.8694296 | 0.06624236 | 0.1095438 | 0.3788617 |
| LOC_Os02g16540 | 0.5916226 | 0.07643837 | 0.7283536 | 0.1013058 | 0.3168398 | 0.7474883 | 0.7041869 | 0.434869 |
| LOC_Os05g50700 | 0.5179651 | 0.8139739 | 0.1620587 | 0.9917181 | 0.5735725 | 0.3620995 | 0.4054009 | 0.6747188 |
| LOC_Os01g46800 | 0.508931 | 0.1591299 | 0.8110451 | 0.01861426 | 0.3995314 | 0.6647967 | 0.6214953 | 0.3521774 |
| LOC_Os03g55080 | 0.3512459 | 0.9806932 | 0.328778 | 0.8415626 | 0.7402918 | 0.1953802 | 0.2386816 | 0.5079995 |
| LOC_Os01g14440 | 0.645524 | 0.6864151 | 0.03449989 | 0.8641592 | 0.4460136 | 0.4896583 | 0.5329597 | 0.8022776 |
| LOC_Os01g53040 | 0.3585379 | 0.309523 | 0.9614382 | 0.1317788 | 0.5499244 | 0.5144036 | 0.4711022 | 0.2017843 |

| | | | | | | | | |
|---|---|---|---|---|---|---|---|---|
| LOC_Os02g08440 | 0.3639203 | 0.9680187 | 0.3161036 | 0.8542371 | 0.7276173 | 0.2080546 | 0.251356 | 0.5206739 |
| LOC_Os01g40260 | 0.521359 | 0.146702 | 0.7986171 | 0.03104222 | 0.3871034 | 0.6772246 | 0.6339232 | 0.3646053 |
| LOC_Os07g40570 | 0.4242032 | 0.2438577 | 0.8957729 | 0.06611353 | 0.4842592 | 0.5800689 | 0.5367675 | 0.2674496 |
| LOC_Os03g33012 | 0.2050332 | 0.4630277 | 0.8850571 | 0.2852835 | 0.7034292 | 0.3608989 | 0.3175975 | 0.04827959 |
| LOC_Os06g44010 | 0.5773865 | 0.09067442 | 0.7425896 | 0.08706978 | 0.3310758 | 0.7332522 | 0.6899508 | 0.4206329 |
| LOC_Os11g45850 | 0.1175517 | 0.5505092 | 0.7975756 | 0.372765 | 0.7909107 | 0.2734174 | 0.230116 | 0.0392019 |
| LOC_Os01g47560 | 0.7545763 | 0.0865153 | 0.5653999 | 0.2642595 | 0.1538861 | 0.9104419 | 0.8671405 | 0.5978226 |
| LOC_Os01g43550 | 0.5879364 | 0.7440026 | 0.09208748 | 0.9217469 | 0.5036013 | 0.4320707 | 0.4753721 | 0.7446901 |
